# ^13^C flux ratio analysis with FRAPPPE reveals differences in metabolic fluxes between gut Bacteroidota and *Escherichia coli*

**DOI:** 10.64898/2026.05.29.728648

**Authors:** Daniel Torka, Bartosz Jan Bartmanski, Alexander Spiegelhalter, Inés Herrera-Gomez, Mariana Barcenas, Bernhard Drotleff, Michael Zimmermann, Maria Zimmermann-Kogadeeva

## Abstract

Gut bacteria shape the metabolism of their host and play an important role in human health. However, systems biology approaches to study their intracellular metabolic fluxes are largely underdeveloped. We present an experimental and computational workflow to quantify metabolic flux ratios in gut bacteria using ^13^C-labeled nutrient supplementation and a newly developed machine learning-based Flux Ratio Prediction Python PackagE (FRAPPPE). We apply FRAPPPE to investigate central carbon metabolism in two prevalent gut Bacteroidota, *Bacteroides uniformis* and *Phocaeicola vulgatus*, in comparison to *Escherichia coli*. FRAPPPE revealed altered tricarboxylic acid cycle bifurcation in Bacteroidota compared to *E. coli* under anaerobic conditions. Further, we used FRAPPPE to investigate co-metabolism of nucleosides and carbohydrates by *B. uniformis* and *P. vulgatus*. We found distinct species-specific patterns in how nucleosides affected growth and were utilized depending on the co-fed compound. We quantified co-metabolism and showed that the tested nucleosides were mainly contributing to anabolic metabolism closely related to the specific co-fed nucleoside. Together, these findings provide insights into central and nucleoside metabolism of two gut Bacteroidota, and showcase FRAPPPE as a generalizable workflow to investigate metabolic fluxes in gut bacteria.

## Introduction

The gut microbiota is a diverse community of microorganisms shaped by metabolic interactions occurring among community members and their host. Such interactions include community members competing for available nutrients and producing metabolic byproducts alongside biomass components and energy. Energy production strategies in the gut microbiota include polysaccharide breakdown, protein digestion (Zeng et al. 2022; Salyers et al. 1977; Yao et al. 2016), and organic acid utilization (Belenguer et al. 2007; Duncan et al. 2002), resulting in various fermentation products dominated by organic acids, such as the short-chain fatty acids acetate, propionate, and butyrate (Mukhopadhya and Louis 2025; Koh et al. 2016). Central carbon metabolism (CCM) consisting of glycolysis, pentose phosphate pathway (PPP) and tricarboxylic acid (TCA) cycle provides both energy and essential biomass building blocks, ultimately orchestrating metabolic activities of a bacterial cell. Identifying which metabolic pathways are used to convert substrates to products and how their activity changes upon perturbations is essential to understand, predict and eventually control metabolic interactions within communities and with the host.

Metabolic pathway activity is usually expressed as metabolic fluxes, or metabolite turnover rates. Metabolic fluxes cannot be directly measured from metabolite or protein abundance data, and have to be inferred using metabolic modeling approaches (Sauer 2006). ^13^C-metabolic flux analysis (^13^C-MFA) (Wiechert 2001) emerged as a key methodology to resolve CCM fluxes. Experimentally, an isotopically labeled carbon source is provided to the culture of interest and the labeling patterns of intracellular metabolites are quantified with mass spectrometry or nuclear magnetic resonance techniques (Long and Antoniewicz 2019). Since carbon atom transitions between individual compounds are reaction-specific, activity of reactions can be inferred from the measured intracellular metabolite labeling patterns. Labeling patterns of neighboring metabolites can be used to estimate flux ratios, e.g. relative contributions of alternative reactions to a metabolite pool, that can be calculated analytically (Fischer and Sauer 2003) or with machine learning workflows (Kogadeeva and Zamboni 2016; Wu et al. 2022). Combined with extracellular metabolite uptake and secretion rates, intracellular metabolite labeling patterns enable the absolute quantification of metabolic fluxes through specific pathways (Rahim et al. 2022; Weitzel et al. 2012; Quek et al. 2009; Foguet et al. 2019).

^13^C-MFA workflows are well established for bacterial model organisms such as *Escherichia coli* and *Bacillus subtilis*, and have been extensively used to investigate CCM activity across different strains, environmental and genetic perturbations (Gonzalez et al. 2017; Crown et al. 2015; He et al. 2014; Okahashi et al. 2017; Rühl et al. 2012; Fischer and Sauer 2005). Their application to non-model organisms, such as gut bacteria, remains challenging. First, it can be challenging to design defined minimal culture media for controlled labelling experiments. Second, the metabolic network structure may be partially unknown, which hampers accurate fitting of the metabolic fluxes based on the labelling data. Previous studies in *Bacteroides fragilis* and *Bacteroides thetaiotaomicron* using ^14^C or ^13^C labelled carbon sources indicated that biosynthesis of succinate, propionate and alpha-ketoglutarate (AKG) in these organisms are different from *E. coli* (Macy et al. 1978; Schofield et al. 2018). Bacteroidota diversity in ability to sustain growth in different minimal media (Tramontano et al. 2018) and on different saccharides (Salyers et al. 1977; Feng et al. 2022) motivates development of systematic and generalized workflows to investigate their CCM and peripheral pathway activity across species and conditions.

We propose an experimental and computational workflow for ^13^C flux ratio analysis tailored to non-model organisms, and demonstrate its applicability by investigating CCM activity in *B. uniformis* and *P. vulgatus* under anaerobic conditions, two of the most abundant and prevalent members of the western gut microbiota (Maier et al. 2018). The key features that ensure flexibility and generalizability of our workflow are the focus on flux ratios and the use of machine learning to build flux ratio predictors trained on simulated datasets. Conceptually, this approach is inspired by the MATLAB-based tool SUMOFLUX (Kogadeeva and Zamboni 2016), which we expand and implement as an open source Flux Ratio Prediction Python PackagE (FRAPPPE). FRAPPPE also enables flux ratio estimation in cases for which exact knowledge of the network structure may be limited and the experimental measurements may be scarce, while offering diagnostic metrics to assess the prediction robustness. Applied to *B. uniformis* and *P. vulgatus* in comparison to *E. coli* grown in glucose minimal medium, FRAPPPE identified differences in the TCA cycle operation that may hint towards different optimization of the TCA cycle. To demonstrate the applicability of FRAPPPE in more complex nutrient conditions, we tested *B. uniformis* and *P. vulgatus* ability to co-utilize nucleosides along with glucose and show that while they do not substantially contribute to energy production, they are used to replenish nucleoside intermediates pools. Taken together, labeling experiments in combination with ^13^C flux ratio analysis with FRAPPPE offer a generalizable platform for the investigation of metabolic fluxes and their adaptation to changing conditions in non-model organisms.

## Methods

### Materials

All chemicals and consumables along with their respective vendors and catalog numbers are also listed in **Supplementary Table S1**.

### Cultivation

Experiments were conducted with three biological replicates unless otherwise specified. All cultivation was conducted in vinyl anaerobic chambers (Coy Laboratory Products Inc.) under anaerobic conditions. The anaerobic atmosphere was based on nitrogen and maintained at < 50 ppm O_2_, 1.5% - 1.8% H_2_, 12% CO_2_. Liquid media was kept for >24 h under anaerobic conditions before usage. For rich culture medium, 41.8 g modified Gifu Anaerobic Medium (mGAM; Nissui Pharmaceutical, 05433) was dissolved in 1 L H_2_O and autoclaved. Glucose minimal medium was based on the defined minimal medium recipe provided in (Bacic and Smith 2008). Mineral 3B solution, 20% glucose solution, hemin solution, FeSO_4_ solution and NaHCO_3_ solution were prepared as previously described (Bacic and Smith 2008). All filter sterilization were performed with a 50 ml syringe and a 0.22 µm filter (Merck Millipore, Ltd., Cat.-No: SLGV033RS). For 160 mM dithiothreitol (DTT) solution, 1.242 g DTT (Roche, Cat.-No: 51280-1G) was dissolved in 50 mL Milli-Q water, filter sterilized and stored at 4 °C. For 160 mM Na_2_S_2_O_3_ solution, 1.266 g Na_2_S_2_O_3_ (Sigma-Aldrich, Cat.-No: 217263-250G) was dissolved in 50 mL Milli-Q water, filter sterilized and stored at 4 °C. For 0.05% Vitamin B_12_ solution, 5 mg Vitamin B_12_ (Sigma-Aldrich, Cat.-No: V2876-250mg) was dissolved in 10 mL Milli-Q water, filter sterilized and stored at 4 °C wrapped in aluminium foil. 1 L glucose minimal medium was prepared by mixing the individual compounds under sterile conditions and used within a day as we observed black precipitate forming after multiple days. The final composition of the minimal medium can be found in **Supplementary Table S2**. For nucleoside co-feeding experiments, glucose minimal medium was supplemented with either 10 mM adenosine (Sigma-Aldrich, A9251), 10 mM cytidine (Sigma-Aldrich, C122106), 10 mM thymidine (TCI, T0233) or 10 mM uridine (Sigma-Aldrich, U3750).

*E. coli* K12 MG1655, *B. uniformis* ATCC 8492 and *P. vulgatus* ATCC 8482 were streaked from cryostocks on mGAM agar plates and incubated under anaerobic conditions for 24 - 72 hours at 37 °C. To prepare starter cultures, a single colony was picked from the agar plate and transferred into 3 mL of mGAM and incubated under anaerobic conditions for 24 h at 37 °C.

### Growth and nutrient uptake quantification under co-feeding conditions

Co-feeding experiments for the quantification of growth and nutrient uptake in glucose and nucleoside minimal medium were conducted with five biological replicates per species and two replicates for bacteria-free medium in 2 mL 96-deep-well plates (Nunc, Thermo Fisher Scientific). Precultures were prepared as described in the previous section; 15 µL of a 1:10 prediluted starter culture (500 µL preculture + 4.5 mL fresh anaerobic minimal medium supplemented with 10 mM glucose) were inoculated in 1.5 mL minimal medium with either 10 mM glucose or 10 mM glucose + 10 mM nucleoside, resulting in a final dilution of 1:1000 in each well. The deep-well plates were sealed with an AeraSeal film (Sigma-Aldrich), and incubated anaerobically at 37 °C. Sampling of the cultures was conducted 0, 12, 15, 17, 22, 24, 36, 40 and 44 hours after inoculation. Per timepoint, 100 uL of culture was transferred into a 96 well round bottom plate and removed from the anaerobic chamber. OD_578_ was measured immediately afterwards using a BioTek Synergy H1 96 well plate reader (Agilent). Areas under the growth curves (AUCs) were calculated using the trapz function of the pracma package (version 2.4.6, https://github.com/cran/pracma) in R. The plate was centrifuged at 3,220 x g for 5 min at 4°C and 90 µl of supernatant were transferred into a fresh round bottom plate. Supernatant plates were sealed with aluminum foil seals (Beckman Coulter Life Sciences) and stored at −80 °C until further analysis.

### Glucose uptake quantification in culture supernatant

Glucose uptake was measured in 1:500-diluted bacterial culture supernatants collected over six time points using the Amplex™ Red Glucose/Glucose Oxidase Assay Kit (Thermo Fisher Scientific, cat. no. A22189) according to the manufacturer’s instructions. Briefly, 50 µL of diluted supernatant was mixed with 50 µL of reaction mix, incubated for 30 min at room temperature protected from light, and fluorescence recorded on a Tecan Spark plate reader (λ_ex_ = 530 nm, λ_em_ = 590 nm). Triplicate calibration curves per plate were prepared from 1:2 serial dilutions of glucose standards (50-3.125 µM), and concentrations were calculated by linear regression, corrected for dilution, and reported as mean ± SD (µM).

### Targeted nucleoside quantification

Absolute quantification of adenosine, cytidine, thymidine, and uridine was performed using hydrophilic interaction liquid chromatography coupled to tandem mass spectrometry (HILIC–LC–MS/MS) in scheduled multiple reaction monitoring (MRM) mode.

Chromatographic separation was carried out at 40 °C on an XBridge BEH Amide XP column (2.1 × 100 mm, 2.5 µm; Waters) using a 1290 Infinity II Bio LC system (Agilent) at a flow rate of 0.35 mL min⁻¹ with 2-µL injections. Mobile phase A consisted of 10 mM ammonium formate with 0.2% (v/v) formic acid and 0.06 mM L-malic acid in water, and mobile phase B consisted of 98% acetonitrile containing identical additives. The gradient was as follows: 96% B was held from 0 to 1.0 min, decreased to 90% B at 1.5 min, 85% B at 2.2 min, and 78% B at 3.0 min, then decreased to 60% B at 3.2 min and held until 3.8 min, followed by re-equilibration to 96% B at 4.0 min, maintained until 6.0 min.

The LC system was coupled to a QTRAP 6500+ mass spectrometer (Sciex) equipped with a Turbo Spray IonDrive source operating in positive ionization mode. Source conditions were: curtain gas: 35 psi; ion source gas 1: 40 psi; ion source gas 2: 60 psi; source temperature: 400 °C; ion spray voltage: 5000 V; and CAD gas set to 8.

Scheduled MRM acquisition was performed at unit mass resolution on Q1 and Q3 using compound-specific retention time windows (±45–60 s), a target cycle time of 600 ms, and dwell times ranging from 91 to 159 ms. For each analyte and its ¹³C-labelled internal standard (U¹³C-adenosine, U¹³C-cytidine, U¹³C-thymidine, U¹³C-uridine), one quantifier and one qualifier transition were monitored. MRM transitions and compound-specific parameters (declustering potentials, collision energies, and cell exit potentials) are provided in **Supplementary Table S3**.

Bacterial culture supernatant samples were diluted 1:500 in 90% (v/v) acetonitrile containing 500 nM of each U-¹³C-labelled internal standard and transferred to V-bottomed 96-well storage plates (Thermo Scientific, AB-1058) and sealed with peelable heat sealing foil sheets (Thermo Scientific, AB-0745) prior to analysis. Pooled quality control (QC) samples were prepared by combining equal aliquots of processed samples to generate representative low, medium, and high concentration levels. Calibration standards for each of the four analytes, spanning a concentration range from 1.22 nM to 20 µM (1:2 serial dilutions), were prepared in the same solvent as the samples and processed identically. Standards and QC samples were stored in glass vials and sealed with caps.

Calibration curves were generated from analyte-to-internal-standard peak area ratios using weighted linear regression (weighting factor as specified per analyte). Data was processed using SCIEX OS software (version 3.4.5.828).

### Fast filtration

Fast filtration protocol of bacterial cultures was applied to collect samples for the measurement of intracellular metabolites, as previously described (Link et al. 2012; Schofield et al. 2018). In brief, minimal medium batch culture was prepared by inoculating 25 mL of glucose minimal medium with 25 µL starter culture in a 100 mL Erlenmeyer flask. To monitor growth of batch cultures, 1 mL of culture was transferred into cuvettes (ratiolab, Cat.-No: 2712120) and optical density at a wavelength of 600 nm was measured with an Ultrospec 10 Cell density meter (Amersham Biosciences). The filtration station consisted of a vacuum pump (KNF Laboport N86 KN.18) attached to a filtration flask (Schott) with glass frit (Millipore, Cat.-No: XX1014702). Washing steps were conducted with Phosphate-Buffered Saline (PBS) without calcium and magnesium deposited with a syringe. 100 mL extraction solution consisted of 40 ml acetonitrile (CHEMSOLUTE, Cat.-No: 2697.2500), 40 mL methanol (CHEMSOLUTE, Cat.-No: 1428.2500) and 20 mL LC-MS-grade H_2_O (CHEMSOLUTE, Cat.- No: 455.2500). Filter papers were managed with a pair of tweezers. After assembly, the filtration station was washed. A 47 mm 0.22 µm PVDF membrane filter (Durapore, Cat.-No: GVWP04700) was placed on top of the filtration setup and washed. Microbial batch culture volume equivalent of 4 mL OD_600_ = 1 or up to 20 mL of culture was deposited onto the filter. For negative controls, either 10 mL minimal medium or H_2_O was deposited onto the filter (n>=2). The filter was washed to remove remaining supernatant and was deposited swiftly into a Petri dish (Greiner Bio-One, Cat.-No: 628102) containing 3 mL of ice-cold extraction solution (Acetonitrile:Methanol:LC-MS-grade H_2_O 2:2:1). Samples were cooled down to - 20 °C for at least 15 min. The sample was evenly divided by transferring 1.3 mL of extract into two 2 mL Eppendorff tubes each. The filter was washed with 1 mL extraction solution, and 500 µL was again deposited into both 2 mL Eppendorff tubes each. The extract was centrifuged for 5 min at 12700 x g at 4 °C and 1.7 mL was transferred into a fresh 2 mL Eppendorff tube. Samples were dried under vacuum at 37 °C using a speed-vac (Genevac EZ-2.4.0 Series Centrifugal Evaporators, Avantor, Inc) until dry and stored at -80 °C until measurement.

### Untargeted LC-MS/MS

Dried samples were dissolved in 100 µL extraction solution (Acetonitrile:Methanol:LC-MS-grade H_2_O 2:2:1). Samples were centrifuged at 21130 x g for 5 min at 4 °C. Supernatant was transferred into glass vials (Macherey Nagel, Cat.-No:702013) with an inlet (Macherey Nagel, Cat.-No: 702077) and sealed with a lid (Macherey Nagel, Cat.-No: 702046). Pooled quality control (QC) samples were prepared by mixing equal aliquots from each processed sample.

LC-MS/MS analysis was performed on a Vanquish UHPLC system coupled to an Orbitrap Exploris 240 high-resolution mass spectrometer (Thermo Fisher Scientific, MA, USA) in negative and positive ESI (electrospray ionization) mode. Chromatographic separation was carried out on an Atlantis Premier BEH Z-HILIC column (Waters, MA, USA; 2.1 mm x 100 mm, 1.7 µm) at a flow rate of 0.25 mL/min. The mobile phase consisted of water:acetonitrile (9:1, v/v; mobile phase phase A) and acetonitrile:water (9:1, v/v; mobile phase B), which were modified with a total buffer concentration of 10 mM ammonium acetate (negative mode) and 10 mM ammonium formate (positive mode), respectively. The aqueous portion of each mobile phase was pH-adjusted (negative mode: pH 9.0 via addition of ammonium hydroxide; positive mode: pH 3.0 via addition of formic acid). The following gradient (20 min total run time including re-equilibration) was applied (time [min]/%B): 0/95, 2/95, 14.5/60, 16/60, 16.5/95, 20/95. Column temperature was maintained at 40 °C, the autosampler was set to 4 °C and sample injection volume was 5 µL. Analytes were recorded via a full scan with a mass resolving power of 120,000 over a mass range from 60 - 900 *m/z* (scan time: 100 ms, RF lens: 70%). To obtain MS/MS fragment spectra, data dependent acquisition was carried out (resolving power: 15,000; scan time: 22 ms; stepped collision energies [%]: 30/50/70; cycle time: 900 ms). Ion source parameters were set to the following values: spray voltage: 4100 V (positive mode) / - 3500 V (negative mode), sheath gas: 30 psi, auxiliary gas: 5 psi, sweep gas: 0 psi, ion transfer tube temperature: 350 °C, vaporizer temperature: 300 °C.

All experimental samples were measured in a randomized manner. Multiple QC samples were injected at the beginning of the analysis for system equilibration, throughout the sequence after every other experimental sample, and at the end to monitor instrument performance over time. An authentic unlabeled sample was measured per batch to account for matrix effects of the sample material and to match MS2 spectra for compound identification. For determination of background signals and subsequent background filtering, additional processed blank samples were recorded. Data was processed using CompoundDiscoverer 3.4.0.1183 (Thermo Fisher Scientific). Level 1 feature identification was based on an in-house library for metabolomics (Dekina et al. 2024) using accurate mass, isotope pattern, MS/MS fragmentation, and retention time information. Level 2 feature annotation was performed with mzCloud (https://www.mzcloud.org/). Features were removed when also identified in water or media blanks, when within a batch the maximum peak quality did not surpass 5, or when more than 70% of samples had a peak quality below 3.5. Resulting isotopologue relative abundance tables were further processed in R 3.4.1.

Multiple feature filtering criteria were applied in the following order. Features were removed if i) their area was below the threefold of the mean media blank area, ii) they did not occur in all three biological replicates, iii) their MS2 spectrum was not found as a “full match” in either the in-house library or mzCloud, iv) their retention time was not within 1 minute tolerance of recorded retention time of the in-house library (only applied to MS2 in-house library matches). We performed the following filtering step on either the entire dataset for the species comparison or separately per species and media condition for co-feeding experiments: Features were removed if any isotopologue standard deviation calculated across biological replicates was > 0.3. Next, we removed features from the species comparison dataset that did not occur in all samples or had a m+0 > 0.25 in any of the samples where 100% [U-^13^C] Glucose was used. Last, we removed features that occurred multiple times (e.g. features measured in both positive and negative mode) by keeping only the feature with the highest peak area. Unfiltered isotopologue data is available in **Supplementary Table S4a and S4b,** filtered datasets are available in **Supplementary Table S4c and S4d** for species comparison and co-feeding experiments, respectively.

### The FRAPPPE workflow

The FRAPPPE workflow was developed in Python 3.10.18. Metabolic models with reactions and atom transitions were built using the FreeFlux software package (version 0.3.6) (Wu et al. 2023). Model reactions for *E. coli* and *B. uniformis/P. vulgatus* models are provided in **Supplementary Tables S5 and S6**. For flux sampling, the hopsy package (version 1.6.1) (Paul et al. 2024) was used to formulate a problem in the form of Ax <= b by implementing steady state constraints alongside upper and lower bounds constraints on the fluxes. The problem was constrained using the stoichiometry matrix derived from the FreeFlux model object. The lower bounds and upper bounds for all fluxes were (0, 100), except for glucose uptake (10,10). In simulations for strains grown under anaerobic conditions, an additional constraint of CO_2_ uptake (90, 100) was formulated. The initialization point for flux sampling was chosen as the Chebyshev center of the convex polytope constraining the sampling space. By default, a Markov chain was instantiated and 100,000 samples were drawn with a thinning parameter of 1,000. After flux sampling, exchange fluxes were constrained by re-sampling of the exchange flux from a uniform distribution between 0 and the net flux value of the reaction for aerobic simulations and the 20-fold of the net flux for anaerobic simulations. For the nucleoside cofeeding condition, three distinct datasets were generated via flux sampling to represent different co-utilisation scenarios: (i) 5,000 flux samples (thinning parameter = 1,000) under baseline conditions, wherein nucleoside uptake was set to zero and flux ratio constraints on CCM were applied; (ii) 5,000 flux samples (thinning parameter = 1,000) wherein nucleoside uptake was constrained to the interval [0, 1] and the flux ratios for nucleoside uptake and phosphorylation were discretized into bins; and (iii) 1,000 flux samples (thinning parameter = 1,000) wherein the dephosphorylation reaction flux was fixed at zero and exchange fluxes were resampled independently in the same manner as the default case. Simulated flux ratios were calculated from simulated fluxes according to the equations in **Supplementary Table S7**. Flux sampling and intracellular metabolite mass distribution vectors (MDV) simulation steps were executed on the EMBL Heidelberg HPC cluster (European Molecular Biology Laboratory *et al*. 2020).

### Flux ratio predictor training

Per ratio, a training data subset was drawn from the original simulated dataset. For the glucose-only case, simulation iterations were binned based on the flux ratio values into 10 distinct bins ranging from (0,0.1), (0.1,0.2), …, (0.9,1). Next, up to 1000 samples were drawn per bin. For the nucleoside cofeeding case, the uptake and phosphorylation flux ratios were each divided into 10 equal intervals, and all 100 pairwise combinations were sampled independently by adding the corresponding bounds directly as constraints in hopsy, drawing 50 samples per bin to ensure uniform coverage across the full ratio space. For these flux subsets, MDVs were simulated for a pre-defined selection of metabolites (**Supplementary Table S8**) using the sim.simulate() function from the FreeFlux package. To represent measurement noise, random values were added to the simulated dataset from the uniform distribution between (−0.01 and 0.01), and the MDVs were subsequently renormalized. Isotopologues for which simulated values were below 0.05 in all simulation iterations were removed before training predictors. The simulated dataset was randomly split into training and test sets with the ratio 2:1 using the train_test_split function of the scikit-learn package (version 1.7.2). Random forest regression models were trained on the training data using the RandomForestRegressor function of the scikit-learn package with its default parameter settings. Model performance was assessed by predicting the flux ratio value on the test set. The mean absolute error (MAE) between true and predicted value on the test set was calculated to evaluate the final model performance.

Several diagnostic plots and metrics were used at each step of FRAPPPE to assess whether the simulated dataset is suitable to analyze the available experimental data. The diagnostic plots include: i) the distributions of all sampled fluxes in the network; ii) the distributions of simulated MDVs overlayed with experimental measurements to check if the simulated data includes experimentally observed values; iii) the histogram of values for the flux ratio of interest (in a general case expected to be uniform between 0 and 1). For flux ratio predictors, the scatterplot depicting expected and predicted flux ratio on the simulated test dataset is reported along with the MAE value. For flux ratio predictions for the experimental data, [0.1 0.9] interquantile range of predictions is reported to assess the robustness of prediction.

### Model formulation

Reactions for central carbon metabolism were derived from the *E. coli* model from (Kogadeeva and Zamboni 2016). The TCA cycle was split into an explicit oxidative and reductive branch. Amino acid reactions were taken from the *E. coli* model from (Wu et al. 2022). A full list of reactions can be found in **Supplementary Tables S5 and S6**. To model nucleoside co-utilization with glucose, the CCM model was extended to include reactions for nucleoside biosynthesis and metabolism. The carbon transitions were taken from the previously published genome-scale model for *E. coli* (Gopalakrishnan and Maranas 2015), where possible. In case the reactions were missing, they were added from the modified genome-scale model of *E. coli* iML1515 (Bernstein et al. 2023), and carbon transitions were checked using the metAMDB (Starke and Wegner 2022) and MetaCyC (Caspi et al. 2020) databases. Models for the adenosine, cytidine, thymidine and uridine co-feeding are provided in **Supplementary table S9a-d**, formulas for the respective flux ratios are provided in **Supplementary table S9e**.

### Statistical analysis

Significance of differences in AUC, glucose and metabolite concentrations were determined using Welch’s t-test (**Supplementary Table S10 and S11c,d**). Significance in difference of labeled carbon share distribution was calculated with a paired Wilcoxon signed-rank test. Correction for multiple testing was conducted with the Benjamini-Hochberg procedure.

## Results

### Establishing a generalized workflow for ^13^C-flux ratio analysis in gut bacteria

We first set out to establish a generalized workflow for flux ratio quantification in gut bacteria focusing on two prevalent commensal bacteria, *Bacteroides uniformis* and *Phocaeicola vulgatus*. As the frequently used Varel-Bryant glucose minimal medium contained methionine and cysteine which in theory could serve as additional carbon sources (Varel and Bryant 1974; Bacic and Smith 2008), we first optimized it to support growth without these additives. We replaced methionine with vitamin B_12_, which is required for methionine biosynthesis in some Bacteroidota and thus enables growth in the absence of methionine (Varel and Bryant 1974) and cysteine with DTT and Na_2_S_2_O_3_, inspired by (Schofield et al. 2018) and (Varel and Bryant 1974) respectively. As intracellular metabolism can rewire within seconds to minutes, keeping the time between sampling and extraction as short as possible is important. The sampling part of our workflow is therefore based on a previously established fast filtration protocol for bacterial batch culture to extract intracellular metabolites (Link et al. 2012; Schofield et al. 2018), and subsequent measurement of labeling patterns using an untargeted LC-MS/MS workflow (Materials and methods).

To ensure that the computational part of the flux ratio analysis workflow is flexible and generally applicable to different bacteria and conditions, we decided to design it based on the surrogate modelling and machine learning workflow proposed in SUMOFLUX (Kogadeeva and Zamboni 2016). We expanded on SUMOFLUX functionalities and implemented our workflow in Python built upon available tools for flux sampling and metabolite labeling pattern simulations, which we called FRAPPPE (Flux Ratio Prediction Python PackagE). The FRAPPPE workflow consists of three main parts (**Figure 1A**): 1) random flux sampling in a given metabolic network using hopsy (Paul et al. 2024); 2) calculation of the mass distribution vectors (MDVs) for intracellular metabolites based on the substrate label using freeflux (Wu et al. 2023); and 3) preparing the dataset and training predictors for flux ratios of interest using random forest regression models from scikit-learn (Pedregosa et al. 2011). To validate our computational workflow, we compared FRAPPPE results step by step with results from SUMOFLUX using simulations and published experimental data for *E. coli* grown aerobically in glucose minimal medium (Kogadeeva and Zamboni 2016; Fischer and Sauer 2003). In the first step, across 10,000 flux sampling iterations, the distributions of sampled fluxes were in good agreement between the two workflows, with the notable exceptions of CO_2_ uptake and release, as well as the reactions pyc (PEP + CO_2_ → OAA) and pck (OAA → PEP + CO_2_), as they are formulated as two independent reactions in the model and thus constrained differently in SUMOFLUX and hopsy flux sampling steps (**Figure S1**). In the second step, using both workflows to calculate MDVs from the same set of flux simulations produced MDVs with high agreement (mean absolute delta in MDV relative abundances of 0.0113) (**Figure S2**). To validate the third step, we first compared the distributions of flux ratios derived from the sampled fluxes used as input to the machine learning training step. The flux ratio distributions in FRAPPPE were less uniform compared to SUMOFLUX distributions, which was improved by binning and subsampling simulation iterations based on the calculated flux ratio before training the regression models. Finally, we built predictors for five flux ratios and applied them to the experimental data for *E. coli* wild type and seven mutant strains with gene deletions in central carbon metabolism reported in (Fischer and Sauer 2003). In four out of five flux ratios, FRAPPPE achieved an equal or better mean absolute error (MAE) for the regressor compared to SUMOFLUX (**Figure 1B**), and provided flux ratios for the experimental data which were in good agreement with the SUMOFLUX estimates (**Figure 1C**). Having established and validated the generalized workflow for ^13^C flux ratio analysis, we next applied it to investigate the central carbon metabolism in gut commensals *B. uniformis* and *P. vulgatus*.

**Figure 1:**
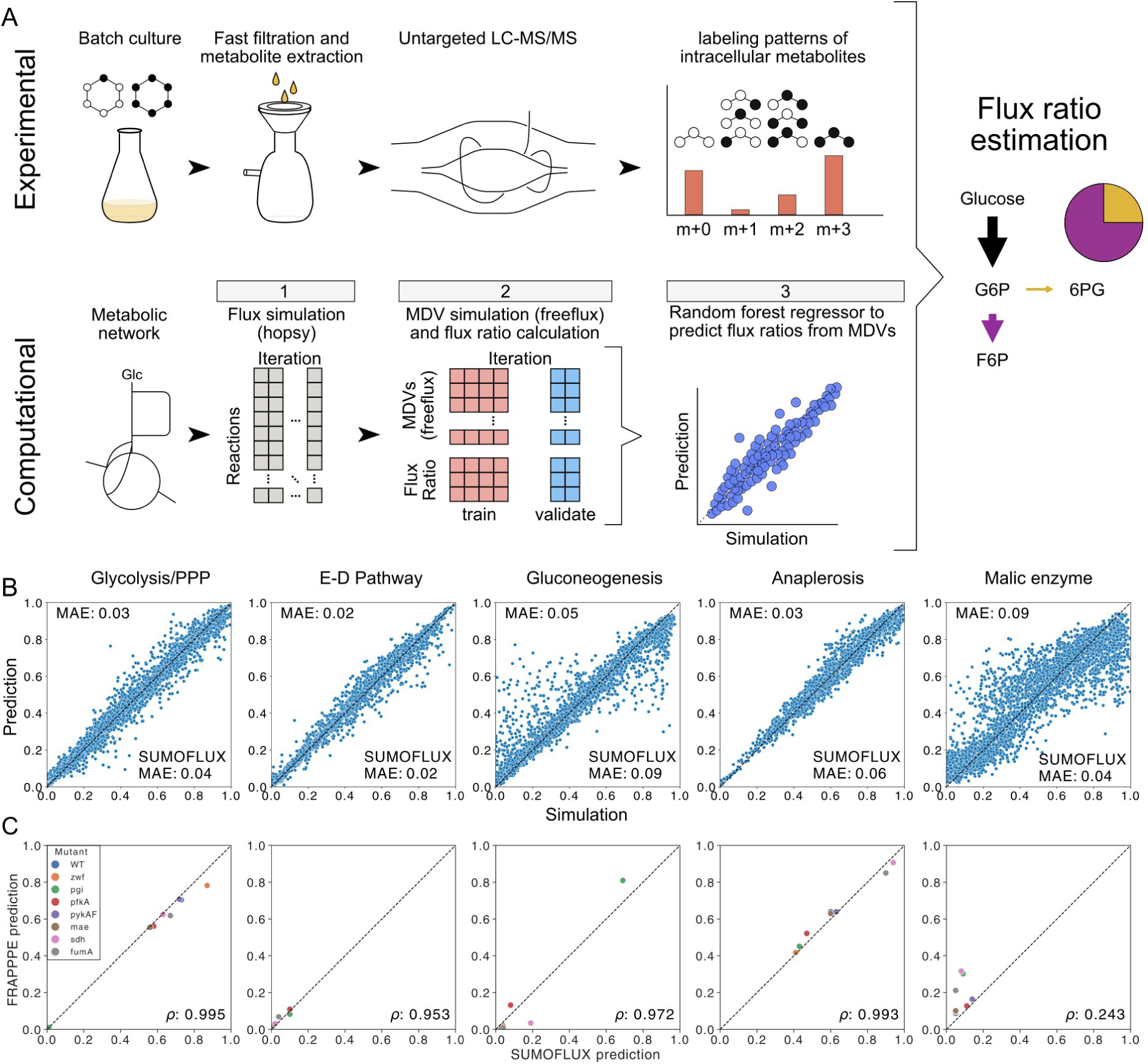
Establishing the FRAPPPE workflow for ^13^C flux ratio quantification. A) Workflow for the quantification of flux ratios based on ^13^C labelling experiments. Top row: Experimental workflow. The species of interest is grown in a batch culture of minimal media with a labeled carbon source. Supernatant is removed via fast filtration, and the intracellular metabolites are extracted from bacteria retained on the filter. Labelled isotopes of intracellular metabolites are measured via untargeted metabolomics. Primary data analysis results in the labeling patterns (MDVs) of intracellular metabolites. Bottom row: Computational workflow with FRAPPPE. A reaction network with carbon transitions is defined as input. Next, flux distributions are sampled. Simulation iterations are subsampled per flux ratio and stratified for flux ratio values to provide a more balanced training dataset. Based on the subsampled flux distributions, flux ratios as well as MDVs of experimentally detected metabolites are calculated. Last, a flux-ratio specific random forest regression model is trained on the subsampled simulated labeling data to predict the flux ratio value from the MDVs. The random forest regressors are applied to the experimental data to predict the flux ratios in the experimental condition. B) Scatter plots of ratio-specific random forest regressor prediction compared to the calculated ratio value from the simulation in the test set. SUMOFLUX MAE values are provided as a reference as reported in (Kogadeeva and Zamboni 2016). C) Flux ratio prediction from real experimental labeling data of *E. coli* with different gene deletions in central carbon metabolism. Comparison between SUMOFLUX and FRAPPPE predictions, ⍴ denotes the pearson pearson correlation calculated between SUMOFLUX and FRAPPPE estimates.

### *B. uniformis* and *P. vulgatus* incorporate more unlabeled carbon in intracellular metabolites than *E. coli*

*B. uniformis* and *P. vulgatus* are among the most abundant and prevalent bacteria of the human gut, yet to our knowledge no previous studies have investigated their central carbon metabolism using ^13^C metabolic flux analysis methodologies. We first performed a baseline characterization of *B. uniformis* and *P. vulgatus* central carbon metabolism and compared it with *E. coli*, a species which has been extensively studied with ^13^C-MFA (Sauer et al. 1999; Crown et al. 2015; Gonzalez et al. 2017). We cultured all three species in the modified Varel-Bryant minimal medium with 10 mM glucose in three different labeling states under strictly anaerobic conditions: 100% [U-^13^C], 100% [1-^13^C] and a mixture of 50% [U-^13^C] and 50% [1-^13^C] in triplicates. We collected samples after 11 hours for *E. coli* and 17 hours for *B. uniformis* and *P. vulgatus* in the mid-to-late exponential growth phase (**Figure 2A**), and quantified intracellular metabolite labeling patterns with an untargeted LC-MS/MS workflow. This workflow, on the one hand, offers accurate identification of metabolites present in the compound library (Materials and methods), while on the other hand can identify differences in labeling patterns of metabolites between species and conditions that were not targeted a priori. In total, we detected labeling patterns of 51 metabolites found in all conditions with the workflow described above (**Supplementary Table 4**).

**Figure 2:**
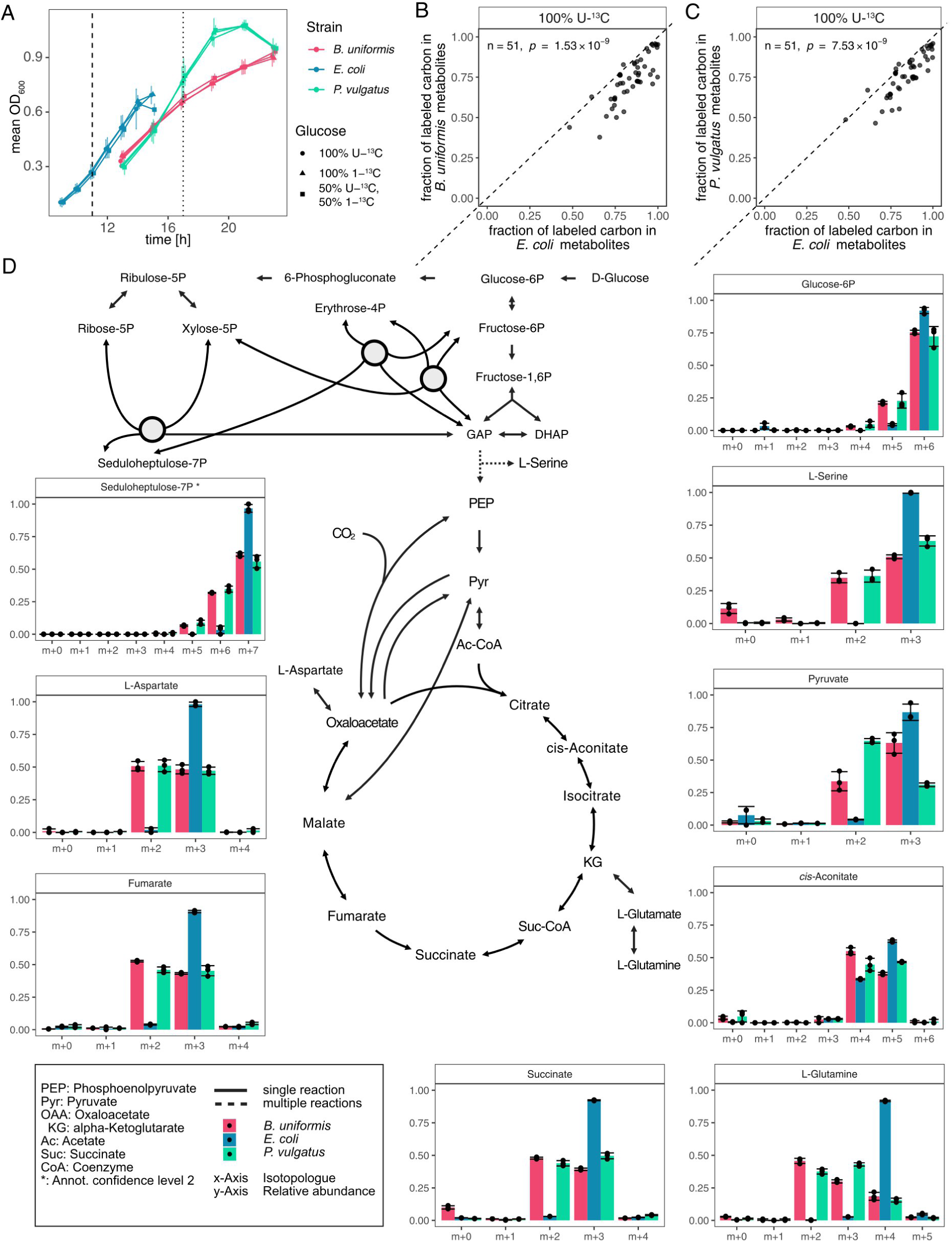
Intracellular metabolites of *B. uniformis* and *P. vulgatus* grown on 100% [U-^13^C] glucose are enriched with unlabeled carbon. A) OD_600_ of batch cultures in different glucose labeling states. Dashed line indicates the start of sampling of *E. coli* cultures, dotted line of *B. uniformis* and *P. vulgatus* cultures. B) Fraction of labeled carbon in intracellular metabolites between *E. coli* and *B. uniformis*. n denotes number of features, *p* statistical significance from a paired Wilcoxon signed-rank test. Each dot represents the mean of three replicates. C) Fraction of labeled carbon in intracellular metabolites between *E. coli* and *P. vulgatus*. n, *p* same as B. D) Center: Scheme of central carbon metabolism including glycolysis, pentose phosphate pathway and TCA cycle. Periphery: MDVs for nine metabolites of the network. Bar height denotes the mean relative abundance across biological replicates, error bars the standard deviation and dots the individual measurements (n=3).

Upon growth on 100% [U-^13^C], the fraction of labeled carbon was significantly lower in *B. uniformis* and *P. vulgatus* compared to *E. coli* metabolites (paired Wilcoxon signed-rank test *p =* 1.15 × 10^-12^ and *p* = 3.61 × 10^-11^, respectively) (**Figure 2B, C**) with an average labeled fraction of 0.635 in *B. uniformis*, 0.678 in *P. vulgatus* and 0.791 in *E. coli*. This effect was less prominent in the 50% [U-^13^C] and 50% [1-^13^C] condition, and not observed in the 100% [1-^13^C] condition (**Figure S3**), likely due to the fact that the majority of carbons in the network are unlabeled and thus indistinguishable between the input carbon sources. We next quantified the cosine similarity of the labeling patterns between the three different species and observed that metabolite labeling patterns in *B. uniformis* and *P. vulgatus* were consistently different to the ones in *E. coli* (**Figure S4**). Visual inspection of metabolite labeling patterns in the 100% [U-^13^C] condition reveals a common pattern. In most *E. coli* metabolites, the MDV is dominated by a single highly abundant isotopologue. Mapped to the central carbon metabolism network, these metabolites can be separated into metabolites upstream of anaplerosis, which remain fully labeled, and metabolites downstream of anaplerosis, which acquired a single unlabeled carbon, derived from the carbonate buffering system and the CO_2_ in the anaerobic atmosphere. In contrast, in most *B. uniformis* and *P. vulgatus* CCM metabolites, more than one highly abundant isotopologue is observed. Already in glucose-6-phosphate, while the most abundant isotopologue is the fully labelled one, more than 20% of molecules in the pool contain one unlabeled carbon. In the TCA cycle intermediates, we observe a more even distribution across two labeling states (either with one or with two unlabeled carbons), while one labeling state (with one unlabeled carbon) is prevalent in *E. coli* (**Figure 2D**). The presence of a single unlabeled carbon can be explained through the incorporation of CO_2_ via anaplerosis, but the origin of the labeling state with two unlabeled carbons remains unclear.

### Flux ratio estimation with FRAPPPE reveals differences in TCA operation between the three species

We next applied the FRAPPPE workflow to investigate which differences in CCM fluxes could explain the observed differences in metabolite labeling patterns. CCM networks of *E. coli* and *B. uniformis* and *P. vulgatus* have several structural differences: while the *E. coli* network contains the Entner-Doudoroff-Pathway, which has not been identified in the two Bacteroidota, the network of *B. uniformis* and *P. vulgatus* has two amphibolic reactions pyruvate carboxylase and oxaloacetate decarboxylase absent in *E. coli*, in addition to phosphoenolpyruvate carboxylase and malic enzyme, which are present in all three species. Further, while *E. coli* converts pyruvate to acetyl-CoA via pyruvate formiate lyase under anaerobic conditions, *B. uniformis* and *P. vulgatus* have genes annotated as ferredoxin-dependent pyruvate oxidoreductase. We therefore constructed two input CCM networks differing in the abovementioned reactions for *E. coli* and *B. uniformis* (which is the same for *P. vulgatus*) (**Supplementary Tables S5 and S6**). We used FRAPPPE to simulate flux distributions and calculate the corresponding MDVs for 15 metabolites in the networks (**Supplementary Table 8**) for the three experimental conditions with differently labeled glucose (100% [U-^13^C], 100% [1-^13^C] and a mixture of 50% [U-^13^C] and 50% [1-^13^C]). We compared simulated MDV distributions to the experimental measurements and confirmed that the experimentally observed labeling states are better represented when the simulations are performed using the corresponding species-specific model (**Figure 3A, B**). Since we were not able to quantify the labeling pattern of citrate, we used the cis-aconitate (intermediate between citrate and isocitrate) MDV in its place. Using these simulated datasets, we next trained flux ratio predictors for a total of 14 flux ratios across both networks and all three labeling states (**Figure 3C, Supplementary Table 7**). We observed that only in 2 out of 13 flux ratios in the *B. uniformis* model predictors achieved MAE <= 0.05. In contrast, in 5 out of 12 flux ratios in the *E. coli* model predictors achieved MAE <= 0.05. For each flux ratio, we selected the best performing predictor based on MAE, and estimated the flux ratio based on the experimental MDVs measured in the corresponding condition (**Figure 3D**). Our predictions based on both the *B. uniformis* and *E. coli* models indicate that glucose is largely utilized through the glycolysis pathway in all three species. This prediction is in agreement with previous descriptions of *E. coli* glucose utilization when cultivated under anaerobic conditions (Sauer et al. 1999; Gonzalez et al. 2017). In the TCA cycle, the predictors highlighted prominent differences between *E. coli* and the two other species: while AKG is produced from citrate through the oxidative branch of the TCA cycle in *E. coli*, it is predominantly produced from succinate through the reductive branch in *B. uniformis* and *P. vulgatus* (**Figure 3D, E**), which is also in agreement with the literature (Allison et al. 1979).

**Figure 3:**
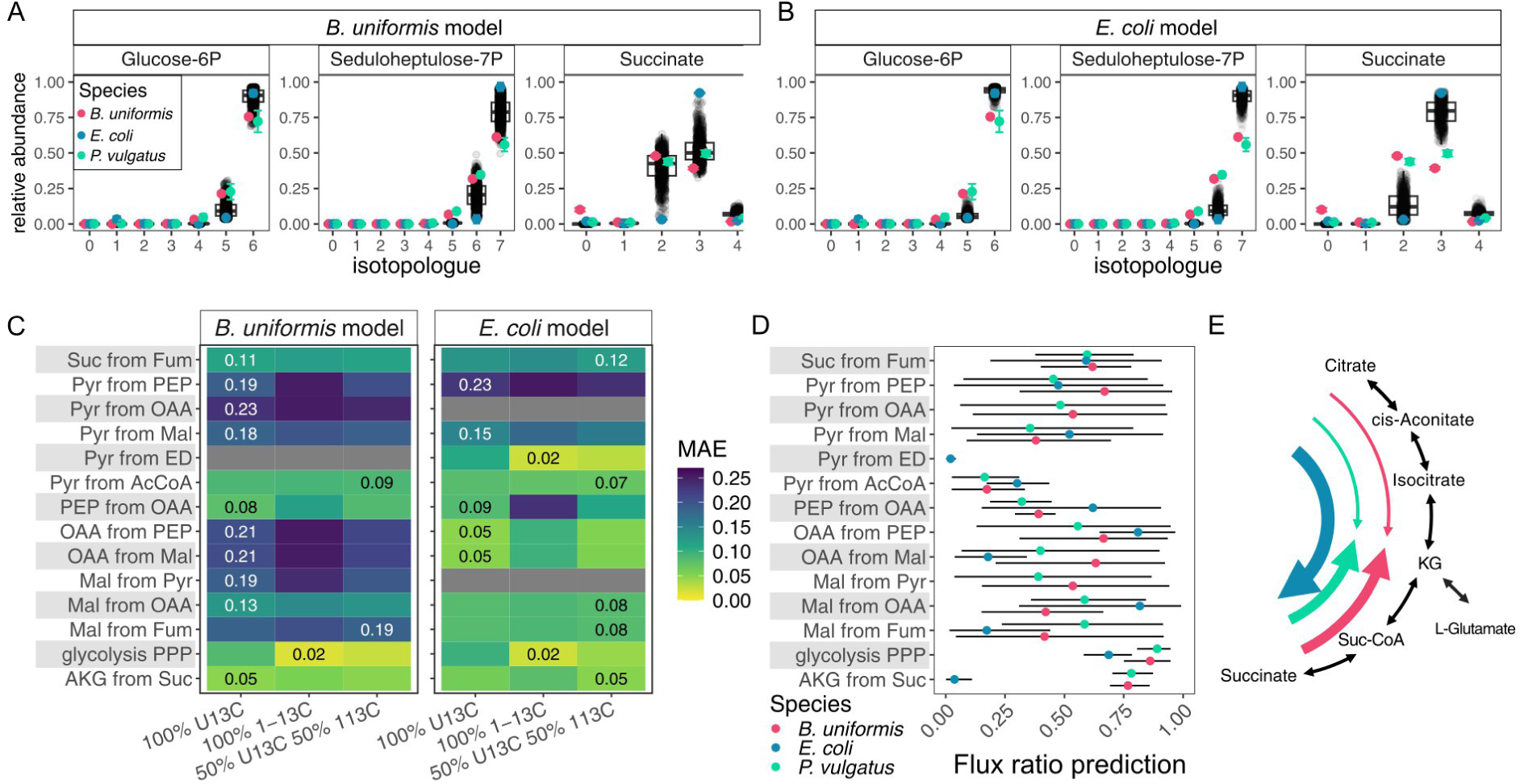
Flux ratio predictions reveal differences in AKG production between *E. coli* and *B. uniformis* and *P. vulgatus*. A) Simulated MDV distribution for 100% [U-^13^C] glucose condition for glucose-6P, seduloheptulose-7P and succinate based on *B. uniformis* model. B) Same as A), based on the *E. coli* model. Colored dots with error bars depict mean and STD of experimental data obtained for each species (n=3). C) FRAPPPE flux ratio predictor performance based on *B. uniformis* model (left) and *E. coli* model (right) assessed as mean absolute error (MAE) on the simulated test set. Best performing ratio predictor indicated by MAE written inside the tile. Grey tiles indicate not predictable as some reactions are not part of the network. D) Flux ratio predictions using the best performing flux ratio predictor and corresponding mean experimental labeling data based on the *B. uniformis* model or *E. coli* model. Bars denote [0.1 0.9] interquantile range of predictions of the random forest predictor. E) Scheme of TCA cycle from citrate to succinate and the FRAPPPE estimates for the flux ratio AKG from Suc. *E. coli* produces ɑKG from citrate, while *B. uniformis* and *P. vulgatus* produce it predominantly from succinate.

### FRAPPPE reveals differences in nucleoside co-metabolism between *B. uniformis* and P. vulgatus

Having established the relative flux distribution in CCM upon growth on glucose, we next investigated how *B. uniformis* and *P. vulgatus* adapt to the presence of more than a single carbon source. We chose to focus on nucleosides, as previous studies showed that supplementation of nucleosides impact community composition (Doo et al. 2017) and have characterized ribose scavenging systems in *B. thetaiotaomicron* (Glowacki et al. 2020). We cultivated *B. uniformis* and *P. vulgatus* with 10 mM [U-^13^C] glucose and 10 mM unlabeled nucleoside (uridine, adenosine, thymidine or cytidine). We observed that supplementation of nucleosides had either positive or negative effect on the growth capacity compared to glucose alone (**Figure 4A**). Adenosine inhibited growth of both species, uridine and cytidine slightly improved growth of both species, although the differences were not significant in *P. vulgatus*, and thymidine had opposite effects: it inhibited *B. uniformis* and promoted growth of *P. vulgatus* (**Figure 4A, S5**, **Supplementary Table S10**). To test the extent of nutrient uptake, we quantified glucose and nucleoside concentrations in sampled supernatants (**Figure S6, Supplementary Table S11**). While we observe a significant reduction in glucose concentration after 17 hours in comparison to 0 hours (two-sided Welch’s t-test, Cytidine cofeeding: 6.94 ± 0.34 mM, *q* = 2.93 × 10^-3^; Thymidine cofeeding: 7.24 ± 0.24 mM, *q* = 1.15 × 10^-3^; Uridine cofeeding: 7.9 ± 0.268 mM, *q* = 1.15 × 10^-3^), only a significant reduction of thymidine could be detected at the latest timepoint in *P. vulgatus* culture (*q* = 1.63 × 10^-3^), indicating that while glucose is the primary carbon source, nucleosides are hardly consumed.

**Figure 4:**
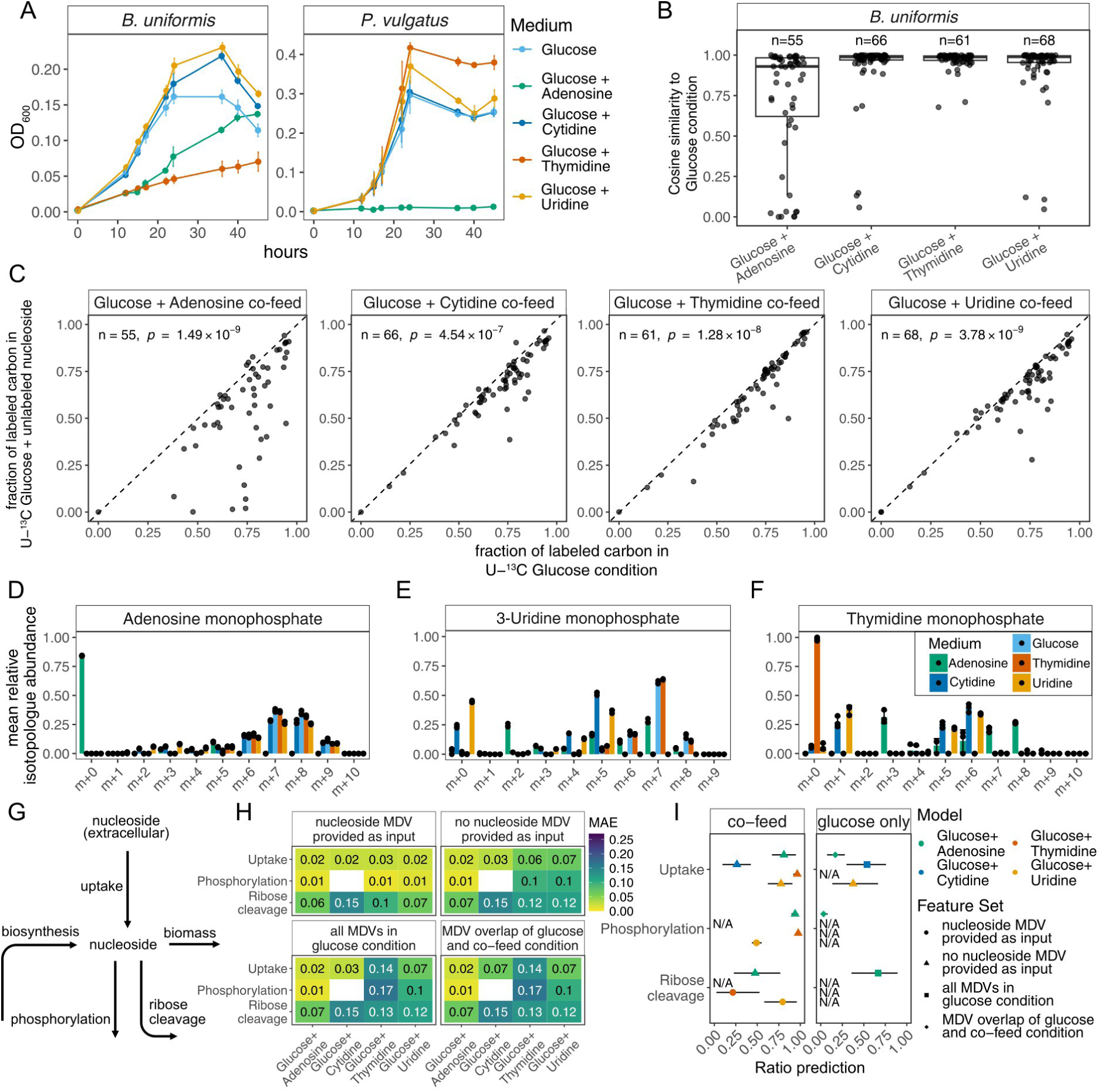
Nucleoside supplementation differentially affects bacterial growth despite their similar metabolism patterns revealed by FRAPPPE. A) Growth of *B. uniformis* and *P. vulgatus* upon nucleoside co-feeding. B) Cosine similarity of *B. uniformis* co-feeding conditions compared to glucose only. C) Fraction of labeled carbon in intracellular metabolites between nucleoside co-feeding condition and glucose alone in *B. uniformis*. n denotes number of features, *p* statistical significance from a paired Wilcoxon signed-rank test. Each dot represents the mean of three replicates. D-F) Labeling patterns of nucleoside intermediates across co-feeding conditions. Bars and whiskers represent mean and STD of n=3 replicates, dots the individual measurements. G) Schematic of nucleoside metabolism reactions used for flux ratio definition. Nucleoside can enter the pool through uptake or biosynthesis, and can either be used for biomass production, phosphorylation or energy production through ribose cleavage. H) Predictor performance heatmap with the labeling pattern of the co-fed nucleoside (top left), without the labeling pattern of the co-fed nucleoside (top right), only with labeling patterns found in the glucose only condition (bottom left) and only labeling patterns found in both the glucose-only and respective co-feeding condition (bottom right). I) Ratio prediction for nucleoside uptake vs de novo synthesis, phosphorylation and ribose cleavage ratios for the experimental data from the respective co-feeding conditions (left) or the glucose only condition (right). Bars denote [0.1 0.9] interquantile range of predictions of the random forest predictor, color denotes the model used for predictions (which in turn defines the respective flux ratio formulations). Shape encodes the model training feature set. N/A indicates cases where no predictor had a MAE < 0.1 on the simulated test set, or the ratio could not be formulated due to the model structure.

To further characterize the extent of nucleoside co-utilization during glucose consumption, we quantified the labeling patterns of intracellular metabolites of *B. uniformis* and *P. vulgatus* and observed a reduction in the cosine similarity of the labeling patterns of several metabolites in comparison to glucose only condition (**Figure 4B, Supplementary Figure S7**). The observed lower number of features confidently identified in adenosine co-feeding condition of *P. vulgatus* is most likely due to poor growth. The fraction of labeled carbon in nucleoside co-feeding conditions in comparison to glucose either remained the same or decreased (**Figure 4C, Supplementary Figure S8**), with the most prominent effect observed in the adenosine co-feeding condition (**Supplementary Table 4**). Multiple nucleoside intermediate and nucleotide labeling patterns were enriched in the [M+0] isotopologue (**Figure 4D-F, Supplementary Figure S9**), indicating that co-fed nucleosides were transported inside the cell and contributed to intracellular intermediate pools. While this pattern was consistent with labeling patterns of nucleosides detected in individual conditions, only minor differences were detected in most of the CCM intermediates and amino acids, indicating that nucleosides are not utilized for energy production to a major extent (**Figure S10, S11**).

To quantify the utilization patterns of the co-fed substrates, we took advantage of the possibility to formulate any flux ratio of interest with FRAPPPE. We first expanded the CCM metabolic models to include nucleoside uptake and metabolism reactions, guided by the metabolites for which the labeling patterns were changing the most compared to glucose only condition (Materials and methods, **Supplementary Table S9**). We then defined three ratios for each co-fed substrate: i) its uptake ratio formulated as a ratio between its uptake versus intracellular production; ii) its phosphorylation ratio formulated as the flux to the corresponding nucleoside phosphate versus all other nucleoside phosphate-producing fluxes; iii) and its cleavage to ribose and nucleotide versus all other consuming fluxes (**Figure 4G**, **Supplementary Table S9**). To investigate the predictors performance dependency on the labeling information on the intracellular nucleoside of interest available as input, we trained predictors on simulated datasets with and without the respective co-fed nucleoside MDV. We observe a gain in performance for many predictors when the nucleoside MDV is added as the input feature (**Figure 4H**). We next applied our trained predictors on the co-feeding data (**Figure 4I**). When not providing the co-fed nucleoside MDV for predictor training, FRAPPPE predictors confidently estimate high relative uptake rates for all nucleosides but cytidine. Further, they estimate that the nucleotides AMP and TMP are synthesized from nucleosides, not from other intermediates. Last, the predictors estimate that the fraction of nucleoside being cleaved into ribose and nucleobase differs widely between co-feeding conditions, however, the overall low predictor performance and large interquantile ranges of predictions for the experimental data indicate that the available measurements are not sufficient to accurately estimate this flux ratio.

## Discussion

Here, we present an experimental and computational workflow for ^13^C flux ratio analysis that is also suitable for applications to non-model organisms and complex conditions. One of its advantages is the use of an untargeted LC-MS/MS platform to quantify metabolite labelling patterns. While it allows confident identification of metabolites of interest commonly used for ^13^C-MFA using in-house and public spectra databases (mzCloud), such as CCM intermediates and amino acids, it can provide a list of candidate metabolites which labeling patterns change between conditions, that can be further investigated and included into the modelling workflow. We use nucleoside co-feeding to showcase how the information from the most differentially labelled metabolites detected by untargeted LC-MS/MS could be incorporated into the network to quantify nucleoside biosynthesis and metabolism flux ratios in *B. uniformis* and *P. vulgatus* with FRAPPPE.

FRAPPPE’s generalizability is ensured by predicting flux ratios with machine learning instead of fitting fluxes to the experimental data, which offers several advantages. First, it allows flexible definitions of the flux ratios of interest, which can be formulated as any combination of the network fluxes to be used as targets for building machine learning predictors. Second, it allows us to identify which of these ratios could be confidently resolved even with scarce measurements, incomplete network information, or complex media with multiple carbon sources. Third, FRAPPPE provides diagnostic metrics and plots that help to assess the distributions of fluxes and flux ratios in the training dataset, the coverage of the experimentally measured MDVs by the simulated ones, and which flux ratios can be accurately predicted from the available data and experimental setup. We demonstrate these advantages by investigating CCM fluxes and nucleoside co-utilization fluxes in gut bacteria. By adjusting the metabolic network reactions and simulating the uptake of unlabeled carbon from CO_2_, we ensure that experimental measurements from *B. uniformis* and *P. vulgatus* are represented in the simulated dataset, which was not the case when *E. coli*-specific model was used for the simulation (**Figure 3A, B**). We further extend the CCM network to include the main parts of nucleoside biosynthesis and metabolism pathways, formulate biosynthesis and metabolism ratios, and assess whether experimental measurements are sufficient to confidently predict these ratios. Adding new reactions to the network however is a challenging task, as one needs to add not only reactions, but carbon transitions as well, which is usually done manually. For *E. coli*, genome-scale ^13^C-MFA models with carbon transitions have been published (Gopalakrishnan and Maranas 2015; He et al. 2016), and carbon transition information across domains of life is provided in databases such as MetAMDB (Starke and Wegner 2022) and MetaCyC (Caspi et al. 2020), offering possibilities for an automated expansion or construction of ^13^C-MFA models for other bacteria in the future. Further, with the acceleration of MDV simulations (Stratmann et al. 2025) and the development of high-throughput experimental workflows for ^13^C-MFA (Nießer et al. 2022; Gurdo et al. 2023) FRAPPPE can be used both for experimental design to select substrate labels and the most informative metabolites to be measured, and for fast flux ratio analysis across thousands of experimental datapoints, if they become available, since the number of samples does not affect the speed of flux ratio estimation by FRAPPPE predictors.

We showcase FRAPPPE by investigating CCM flux ratios in two prominent gut bacteria, *B. uniformis* and *P. vulgatus*, in comparison with a well studied model organism *E. coli*. Using our experimental setup and available measurements, we could estimate two tested flux ratios with very high confidence (MAE<0.05): glycolysis versus PPP and alpha-ketoglutarate from succinate versus citrate. While in all three species the main glucose utilization route was glycolysis, the fluxes around the alpha-ketoglutarate node substantially differed, with *E. coli* producing it from the oxidative branch, and the two *Bacteroidota* from the reductive branch of the TCA cycle. While several other flux ratio predictors had good accuracy (MAE<0.10), they could not confidently identify differences in flux ratios between the three species that would explain large differences in the labelling patterns of the TCA cycle intermediates. Anaplerosis as the sole means of carbon assimilation explains the labeling patterns observed in *E. coli* (oxaloacetate being largely produced from pyruvate). High anaplerotic flux is also consistent with labelling patterns of oxaloacetate and other TCA cycle intermediates in *B. uniformis* and *P. vulgatus,* which have a high fraction of isotopologues with two unlabeled carbons, since pyruvate label shows similar pattern. However, how the pyruvate [M+2] isotopologue is produced could not be resolved with our labelling conditions and measurements. The fact that the growth capacity of multiple species from the gut microbiome is sensitive to bicarbonate concentration (Franke and Deppenmeier 2018) might indicate that CO_2_ assimilation contributes to the high levels of [M+2] in pyruvate and TCA intermediates. Since higher fractions of less labeled isotopologues are already observed in glycolysis intermediates, the incorporation of unlabeled carbon could also happen due to high reshuffling in the transketolase reactions in the PPP. However, we cannot exclude that dithiothreitol, which we provide as a reducing agent to our minimal medium, contributes to the total unlabeled carbon pool, although *E. coli* does not show any signs of assimilation of unlabeled carbon in the same condition. In case the additional unlabeled carbon originates from CO_2_, either anaplerotic products are circulated to a greater extent than previously thought, or additional mechanisms through which CO_2_ is assimilated are at play.

We demonstrated how the FRAPPPE workflow can be extended to more complex nutrient conditions by investigating nucleoside co-utilization with glucose in the two Bacteroidota species. Ribose scavenging systems for the utilization of ribose from nucleosides have been identified across multiple Bacteroidota species (Glowacki et al. 2020), yet the extent to which such riboses contribute to central carbon metabolism in presence of glucose has not been investigated. When provided with equimolar amounts of adenosine and glucose, *B. uniformis* and *P. vulgatus* both showcased a severe reduction in growth. Previous reports show growth delay in both *E. coli* and *Salmonella enterica* upon supplementation with adenosine (Kao et al. 2017; Kitzenberg et al. 2022). Whether physiologically relevant concentrations of adenosine have a growth-inhibiting effect on *B. uniformis* and *P. vulgatus* still needs to be validated. Interestingly, while adenosine shows a consistent growth inhibitory effect in both species, supplementation of thymidine only inhibits *B. uniformis* growth. Double thymidine block is a well-established method to synchronise cell cycle in various cell types by inhibiting the synthesis of dCDP (Bostock 1971; Chen and Deng 2018), but whether this exact mechanism is also inhibiting growth of *B. uniformis*, and why this does not inhibit *P. vulgatus* growth, remains to be investigated. Despite providing various alternative potential carbon sources in the form of nucleosides, labeling patterns in downstream TCA cycle metabolites remained largely unaffected, which indicates a clear preference of glucose over ribose from nucleosides.

Overall, our work provides a scalable and generalizable workflow for ^13^C metabolic flux ratio analysis, and sheds light on the specificities of CCM operation and nucleoside utilization in two prominent gut bacteria, *B. uniformis* and *P. vulgatus*. FRAPPPE might become a valuable building block for the analysis of high-throughput labeling experiments as machine learning predictors, once trained, can be applied iteratively across experimental settings at low computational cost. In combination with high-throughput bacterial cultivation techniques applied to more non-model organisms, we envision that FRAPPPE will become a valuable tool to generate novel insights into their metabolism across species and conditions in the future.

## Data and code availability statement

FRAPPPE source code, tutorials, and example notebooks are freely available at https://github.com/zimmermann-kogadeeva-group/FRAPPPE. Code and data required to reproduce results from the study are available at https://github.com/zimmermann-kogadeeva-group/13CFluxRatioAnalyisWithFRAPPPE.

Preprocessed data and tables used in the study are available on Zenodo: https://zenodo.org/records/20395585.

Raw metabolomics data is available in Metabolights database with accession number REQ20260430219182 at https://www.ebi.ac.uk/metabolights/editor/REQ20260430219182/overview.

## Author contributions

D.B.T. and M.Z.-K. conceptualized the overall project and goals. D.B.T. and B.J.B. developed FRAPPPE, and wrote all scripts used to analyse data and create figures. D.B.T., A.S. and I.H.-G. performed the experiments. B.D. and A.S. developed the metabolomics method to measure nucleosides. D.B.T., M.B., B.D. and A.S. performed metabolomics measurements and data analysis. M.Z. supervised metabolomics measurements and provided experimental advice. D.B.T. and M.Z.-K. wrote the first draft of the manuscript, and all authors read, edited, and approved the final manuscript.

## Acknowledgements

We would like to thank members of the Zimmermann-Kogadeeva lab for helpful discussions and feedback. We also thank the EMBL Metabolomics core facility for the support with metabolomics analysis, the EMBL Microbial Automation and Culturomics core facility for access to laboratory equipment, and the EMBL IT Services staff for managing and provision of access to the HPC resources.

## Declaration of generative AI use

In the preparation of this research paper, we utilized generative AI tools (ChatGPT 5.1 and perplexity.ai) to enhance the clarity and coherence of our writing. We carefully reviewed and edited the AI-generated recommendations to maintain the integrity and authenticity of our work. Generative AI tools were not used to produce new content, but only to rephrase human-written text to improve its clarity. ChatGPT 5.1 and Claude Sonnet were also used for code improvement and debugging, which was subsequently edited by the authors.

## Supplementary Tables

**Supplementary Table S1.** List of strains, chemicals and other materials used in the study.

**Supplementary Table S2.** Minimal medium composition.

**Supplementary Table S3.** MRM details for the quantification of nucleosides with hydrophilic interaction liquid chromatography coupled to tandem mass spectrometry (HILIC–LC–MS/MS).

**Supplementary Table 4.** (A) Unfiltered metabolomics areas and labeling pattern measurements of species comparison experiment in glucose minimal medium condition. (B) Unfiltered metabolomics areas and labeling pattern measurements from glucose + nucleoside co-feeding experiments. (C) Filtered metabolomics areas and labeling pattern measurements of species comparison experiment in glucose minimal medium condition. (D) Filtered metabolomics areas and labeling pattern measurements from glucose + nucleoside co-feeding experiments.

**Supplementary Table S5.** Metabolic network of *E. coli* under anaerobic conditions.

**Supplementary Table S6.** Metabolic network of *B. uniformis* and *P. vulgatus* under anaerobic conditions.

**Supplementary Table S7.** Flux ratio formulations for *E. coli* model and *B. uniformis* models.

**Supplementary Table S8.** List of metabolite MDVs used for training the FRAPPPE prediction models.

**Supplementary Table S9.** (A) Metabolic model for *B. uniformis* glucose + adenosine co-feeding condition. (B) Metabolic model for *B. uniformis* glucose + cytidine co-feeding condition. (C) Metabolic model for *B. uniformis* glucose + thymidine co-feeding condition. (D) Metabolic model for *B. uniformis* glucose + uridine co-feeding condition. (E) Flux ratios formulated for nucleoside co-feeding conditions.

**Supplementary Table S10.** AUC means and significance estimates for *B. uniformis* and *P. vulgatus* growth upon nucleoside supplementation to the glucose minimal medium.

**Supplementary Table S11.** (A) Glucose concentrations of *B. uniformis* and *P. vulgatus* supernatant. (B) Nucleoside concentrations of *B. uniformis* and *P. vulgatus* supernatant. (C) Statistical comparison of glucose concentrations across time points. (D) Statistical comparison of nucleoside concentrations across time points.

## Supplementary Figures

**Figure S1:**
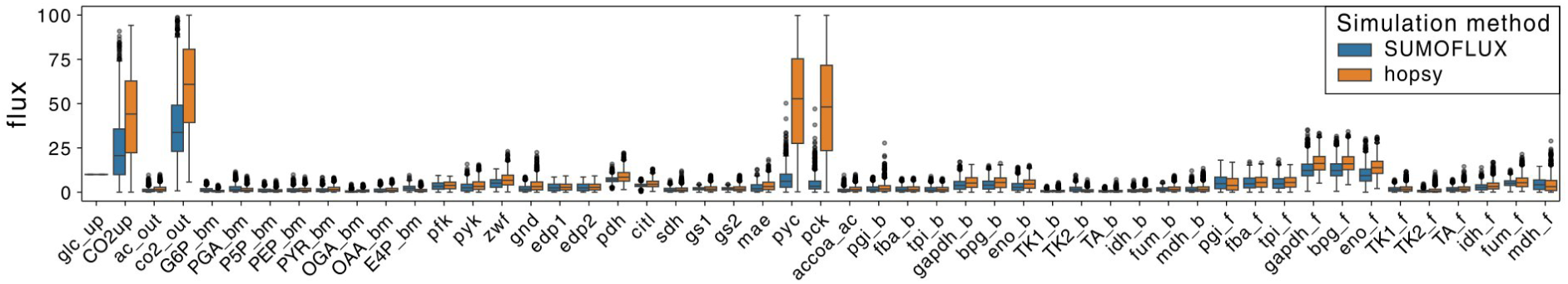
Simulated flux distribution (10000 simulation events) of SUMOFLUX and hopsy with glucose uptake (glc_up) constrained to [10 10]. Reversible reactions are simulated as forward and reverse flux denoted with _f and _b, respectively.

**Figure S2:**
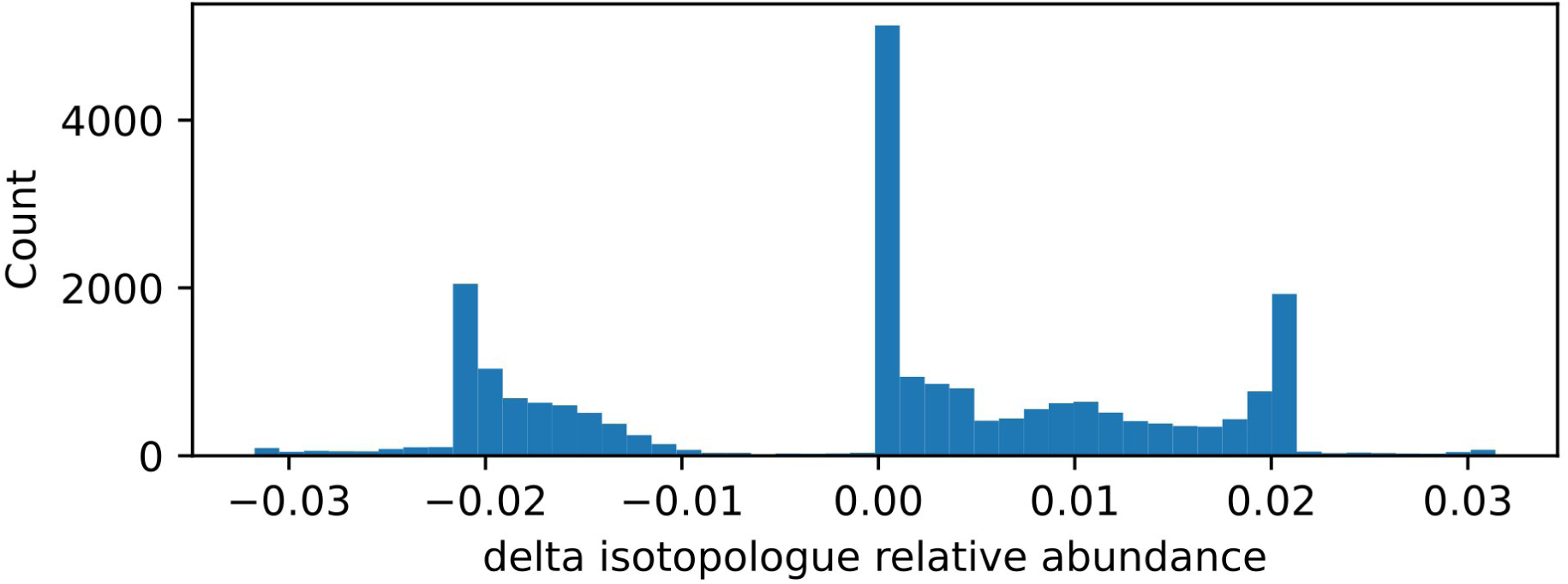
Differences in MDV relative abundances calculated from SUMOFLUX and FreeFlux using the same simulated flux dataset (1000 simulation iterations). The mean absolute difference between the two MDV relative abundance distributions is 0.0114.

**Figure S3:**
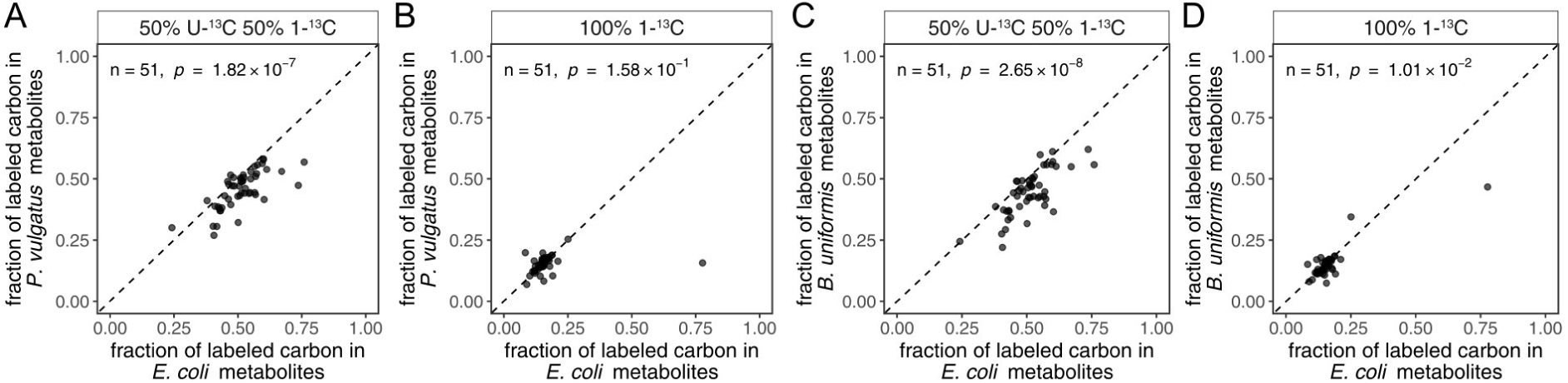
Fraction of labeled carbon in intracellular metabolites in *E. coli* and *B. uniformis* (A,B) and *P. vulgatus* (C,D). While in a mixture of U-^13^C-Glucose and 1-^13^C-Glucose in both Bacteroidota species still display an overall reduction in unlabeled carbon compared to *E. coli*, this effect is not present in the 1-^13^C-Glucose condition. n denotes number of features, p statistical significance from a paired Wilcoxon signed-rank test. Each dot represents the mean of three replicates.

**Figure S4:**
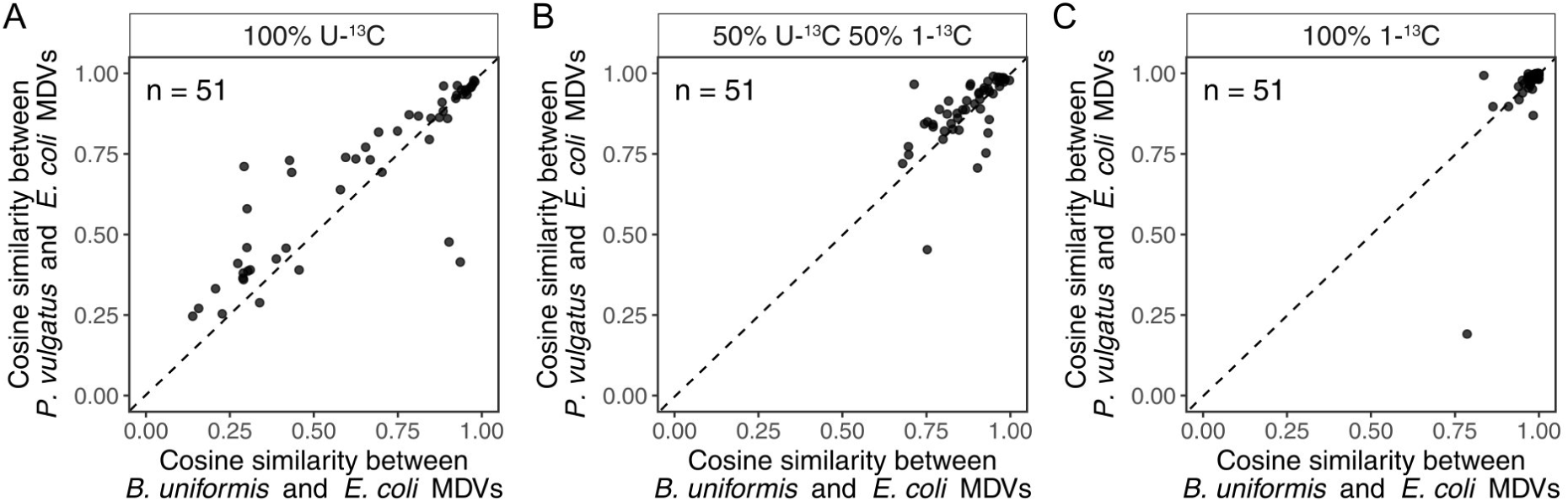
Cosine similarity between metabolite MDVs between *E. coli* and *B. uniformis* against *E. coli* and *P. vulgatus*. In 100% U-^13^C are large discrepancies in labeling patterns which are consistent between in both comparisons (close to the diagonal). A mixture of U-^13^C-Glucose and 1-^13^C-Glucose as well as 1-^13^C-Glucose alone show smaller differences. n denotes number of features, each dot represents the mean of three replicates.

**Figure S5:**
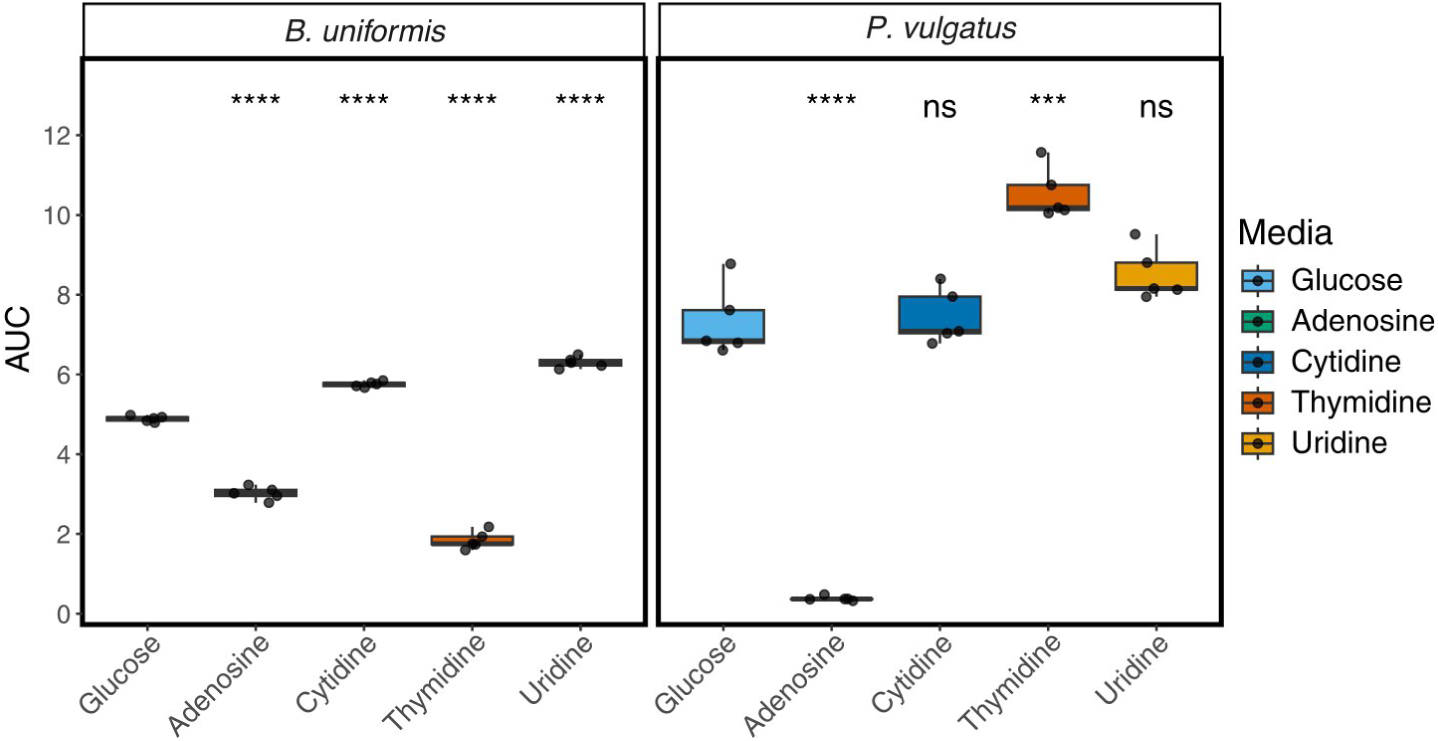
Areas under the curves (AUCs) from growth curves measured under different co-feeding conditions. While we observe significant reduction or increase of growth in all *B. uniformis* co-feed conditions compared to growth on glucose alone, we only observe significant changes in adenosine and thymidine co-feeding conditions in *P. vulgatus* compared to growth on glucose alone. Significance levels: ns: p-value > 0.05, *: p-value < 0.05, **: p-value < 0.01, ***: p-value < 0.001,****: p-value < 0.0001. P-values were calculated using Welch’s t-test by comparing each nucleoside co-feeding condition to glucose alone from n=5 replicates.

**Figure S6:**
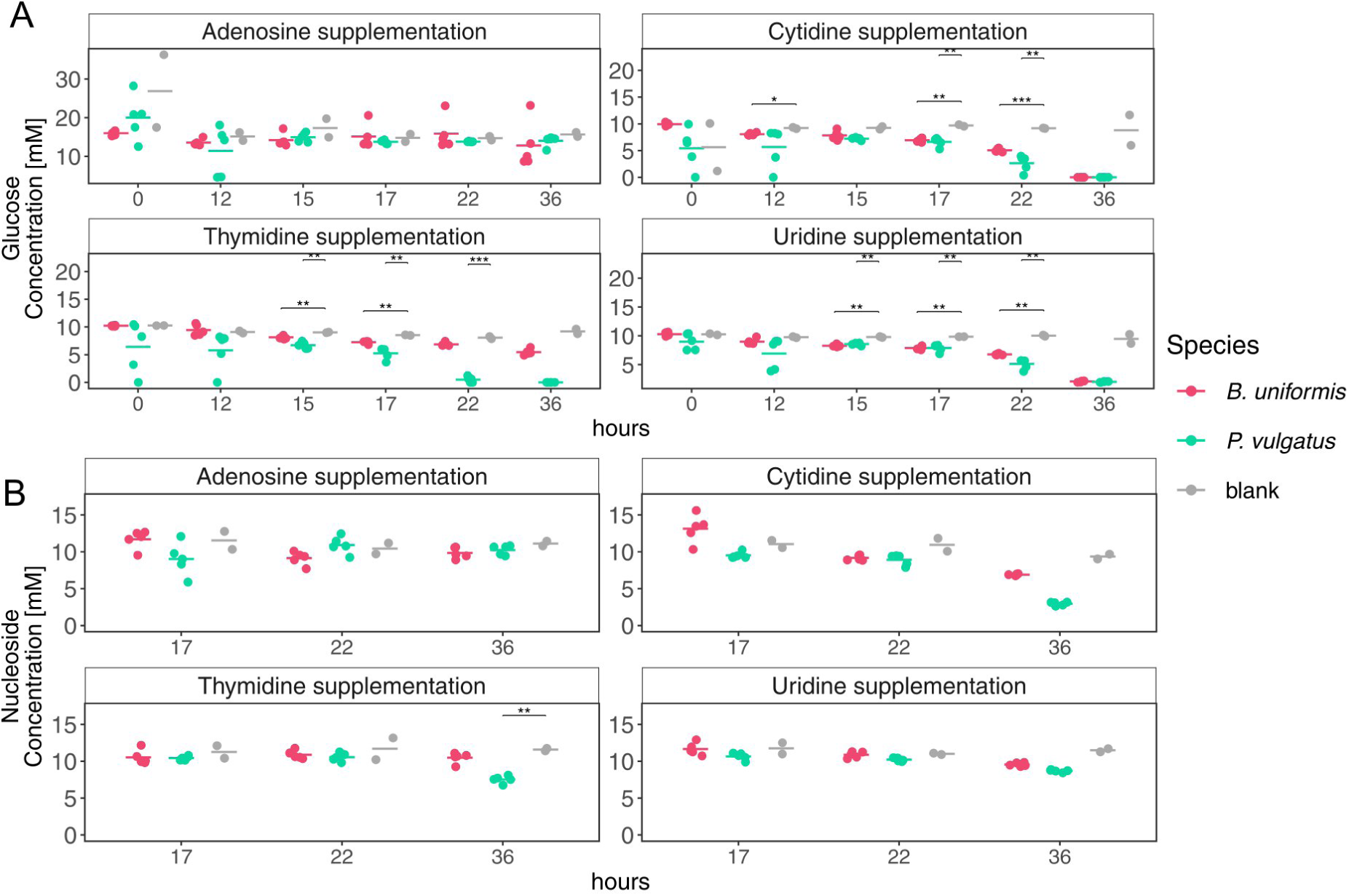
Quantification of glucose and nucleoside uptake in *B. uniformis* and *P. vulgatus* A) Glucose concentrations in supernatant of *B. uniformis*, *P. vulgatus* and bacteria-free media controls (blank) in nucleoside co-feeding conditions. B) Nucleoside concentrations in supernatant of *B. uniformis*, *P. vulgatus* and bacteria-free media controls (blank) in nucleoside co-feeding conditions. Dots represent the individual sample values, the vertical lines denote the groups mean concentration (n=5 for bacterial samples, n=2 for blanks).

**Figure S7.**
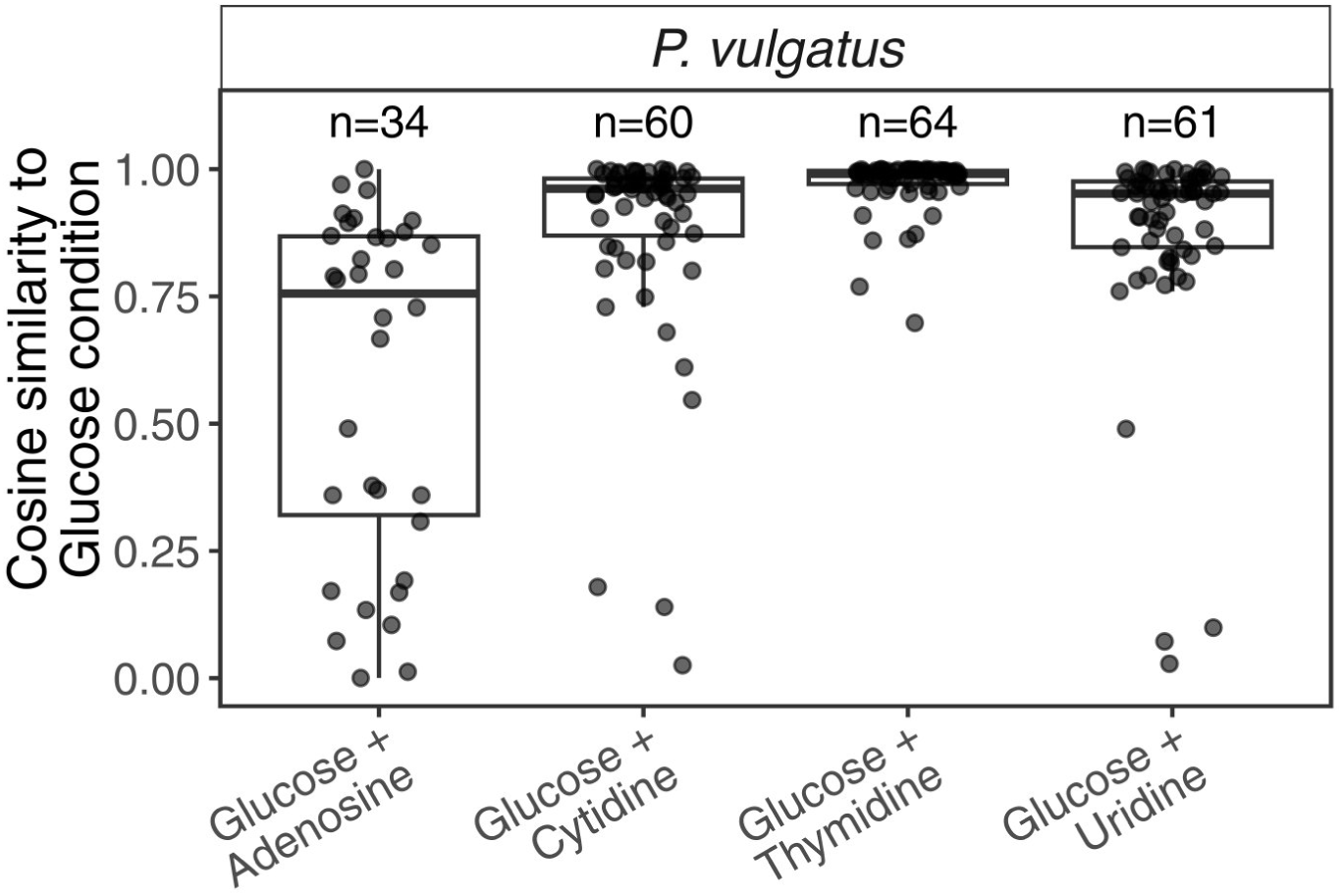
Cosine similarity of *P. vulgatus* co-feeding conditions compared to glucose only. n denotes number of features, each dot represents the mean of three replicates.

**Figure S8.**
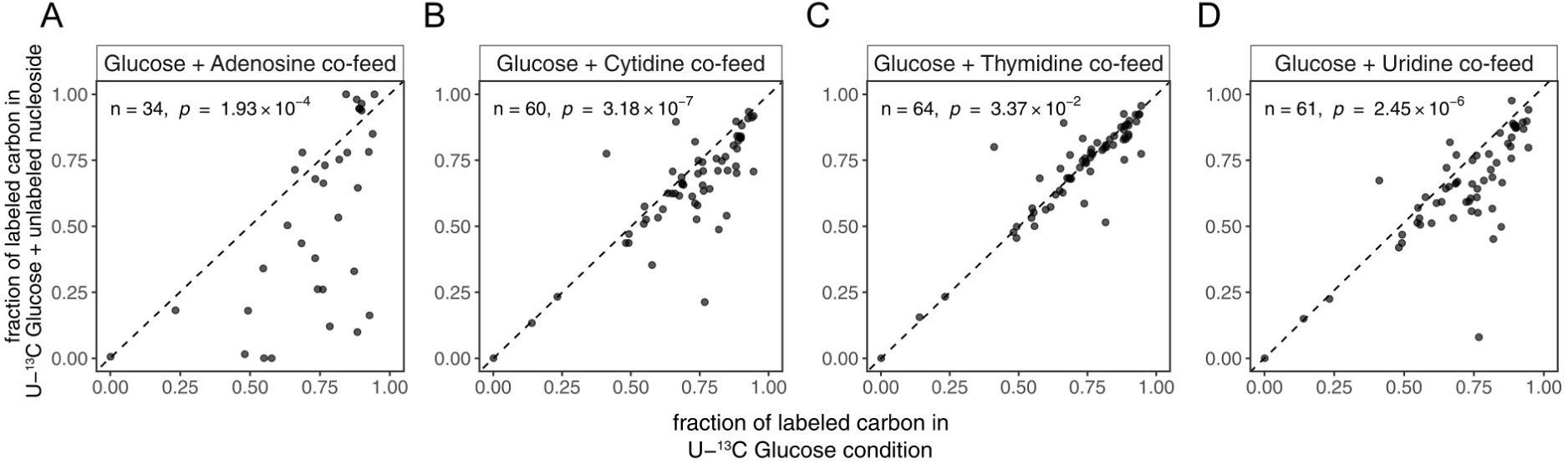
Fraction of labeled carbon in intracellular metabolites between nucleoside co-feeding condition and glucose alone in *P. vulgatus*. n denotes number of features, p statistical significance from a paired Wilcoxon signed-rank test. Each dot represents the mean of three replicates.

**Figure S9:**
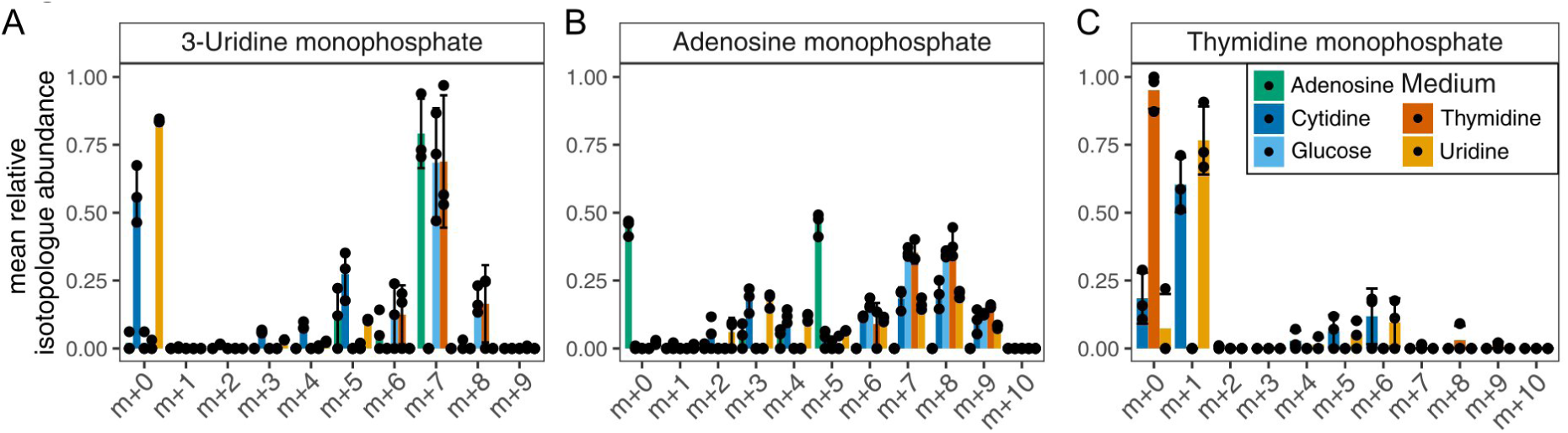
Labeling patterns of nucleoside intermediates in *P. vulgatus* across co-feeding conditions. Bars and whiskers represent mean and STD of n=3 replicates, dots represent the individual measurements.

**Figure S10.**
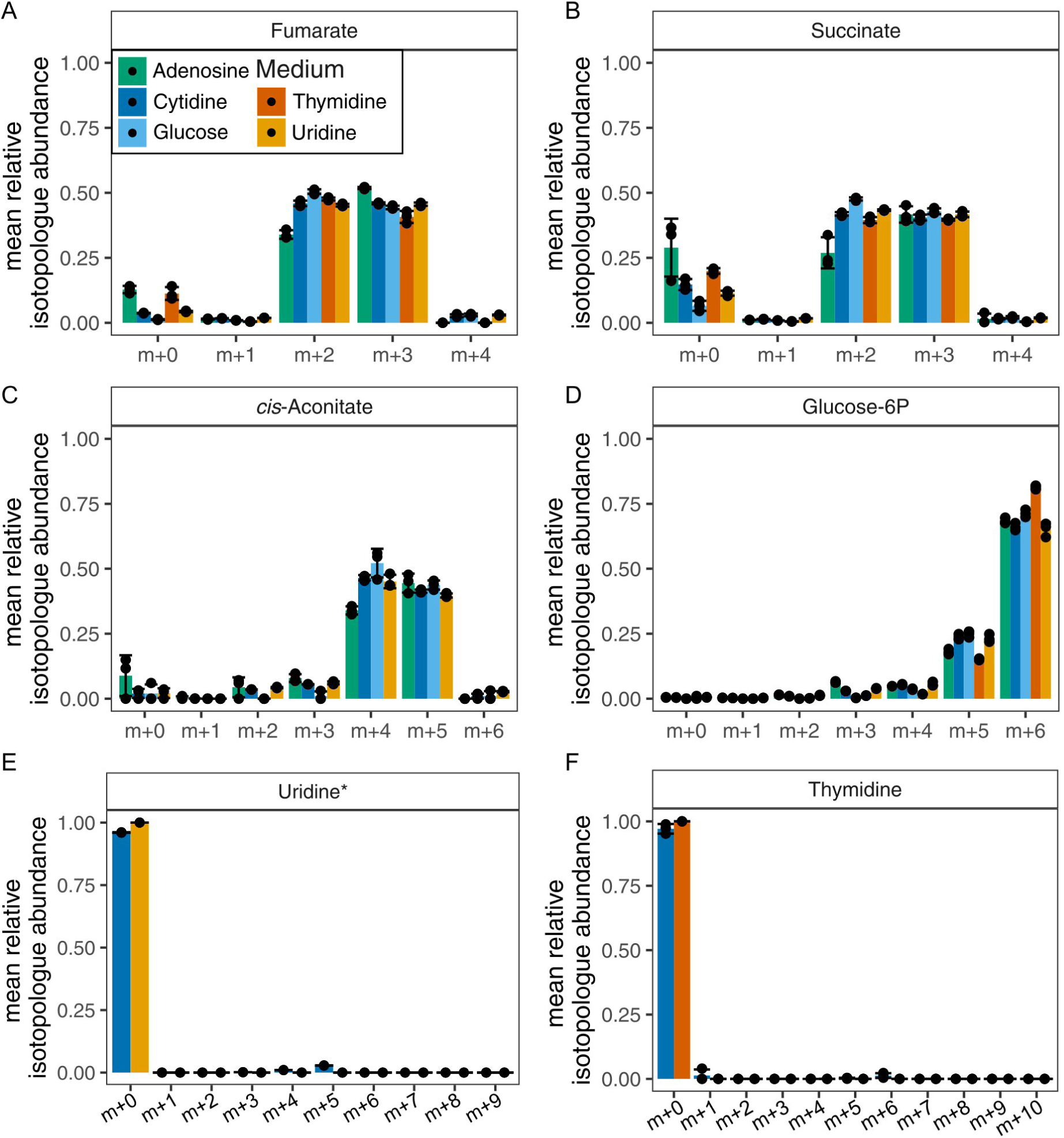
Labeling patterns of nucleosides and central carbon metabolism compounds in Nucleoside co-feeding of *B. uniformis*. Labeling patterns of nucleosides and CCM compounds in U-^13^C glucose and unlabeled nucleoside co-feeding conditions of *B. uniformis*. The observed labeling patterns of CCM metabolites do not show major signs of enrichment for unlabeled carbon in comparison to the glucose-only condition, while nucleosides are enriched for m+0 in few conditions, but not observed consistently in all conditions. Bars and whiskers represent mean and STD of n=3 replicates, dots the individual measurements. * indicates level 2 confidence annotation.

**Figure S11.**
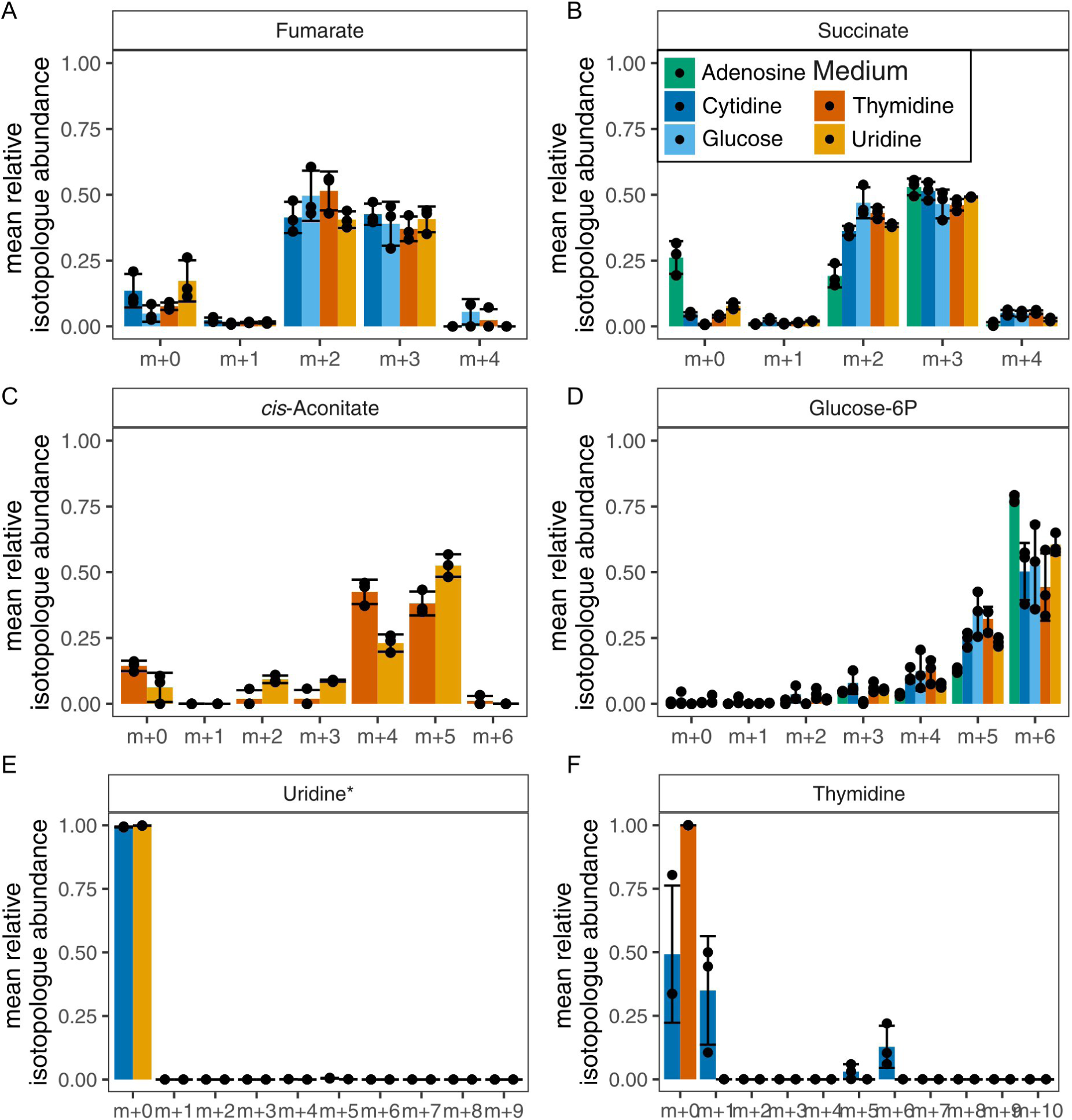
Labeling patterns of nucleosides and central carbon metabolism compounds in Nucleoside co-feeding of *P. vulgatus*. Labeling patterns of nucleosides and CCM compounds in U-^13^C glucose and unlabeled nucleoside co-feeding conditions of *P. vulgatus*. The observed labeling patterns of CCM metabolites do not show major signs of enrichment for unlabeled carbon in comparison to the glucose-only condition, while some nucleosides are enriched for m+0 in few conditions, but not observed consistently in all conditions. Bars and whiskers represent mean and STD of n=3 replicates, dots the individual measurements. * indicates level 2 confidence annotation.

